# Cell density-dependent ferroptosis in breast cancer is induced by accumulation of polyunsaturated fatty acid-enriched triacylglycerides

**DOI:** 10.1101/417949

**Authors:** Elena Panzilius, Felix Holstein, Jonas Dehairs, Mélanie Planque, Christine von Toerne, Ann-Christine Koenig, Sebastian Doll, Marie Bannier-Hélaouët, Hilary M. Ganz, Stefanie M. Hauck, Ali Talebi, Johannes V. Swinnen, Sarah-Maria Fendt, José P. Friedmann Angeli, Marcus Conrad, Christina H. Scheel

## Abstract

Ferroptosis is a regulated form of necrotic cell death caused by iron-dependent phospholipid peroxidation. It can be induced by inhibiting glutathione peroxidase 4 (GPX4), the key enzyme for efficiently reducing peroxides within phospholipid bilayers. Recent data suggest that cancer cells undergoing EMT (dedifferentiation) and those resistant to standard therapy expose a high vulnerability toward ferroptosis. Although recent studies have begun to identify and characterize the metabolic and genetic determinants underlying ferroptosis, many mechanisms that dictate ferroptosis sensitivity remain unknown. Here, we show that low cell density sensitizes primary mammary epithelial and breast cancer cells to ferroptosis induced by GPX4 inhibition, whereas high cell density confers resistance. These effects occur irrespective of oncogenic signaling, cellular phenotype and expression of the fatty acid ligase acyl-CoA synthetase long chain family member 4 (ACSL4). By contrast, we show that a massive accumulation of neutral triacylglycerides (TAG) enriched with polyunsaturated fatty acids (PUFA) is induced at low cell density. In addition, *de novo* lipogenesis and desaturation pathways were found to be reduced at low cell density, indicative of increased fatty acid uptake. Our study suggests that PUFA-mediated toxicity is limited by the enrichment in TAGs that in turn might pose a vulnerability towards ferroptosis. Conclusively, cell density regulates lipid metabolism of breast epithelial and cancer cells, which results in a ferroptosis-sensitive cell state with the potential to be exploited therapeutically during metastatic dissemination.

## Introduction

Ferroptosis, a distinctive form of regulated necrosis ^1, 2^, is emerging as one of the central cell death mechanisms accounting for early cell loss and organ dysfunction in degenerative disease ^3^. On the contrary, induction of ferroptosis may unleash still untapped and highly promising nodes for therapeutic intervention in specific cancer entities ^4^. Accumulating evidence suggests that therapy-resistant cancer cells and those undergoing epithelial-mesenchymal transition (EMT) may become highly sensitive to ferroptosis activation ^5–7^.

Ferroptosis is caused by the accumulation of lethal amounts of peroxidized phospholipids and in this context, it can be triggered by pharmacological inhibition or genetic deletion of glutathione peroxidase 4 (GPX4), the key enzyme directly reducing phospholipid hydroperoxides (PLOOH) to their corresponding alcohols in membranes ^8–10^. As its name implies, ferroptosis is dependent on intracellular iron, which is required for triggering autoxidation of PUFA residues in lipid bilayers ^1^. Moreover, availability of cyst(e)ine inevitably affects glutathione (GSH) metabolism ^11^, and thus GPX4 activity and ferroptosis ^1, 8, 12^. Besides amino acid metabolism, lipid metabolism emerges as one of the key determinants of ferroptosis sensitivity. In this context, the fatty acid ligase acyl-CoA synthetase long chain family member 4 (ACSL4) has been shown to sensitize cells to ferroptosis by activating long-chain poly-unsaturated fatty acids (PUFAs) through formation of PUFA acyl-CoAs (PUFA-CoA) ^13, 14^. Activated PUFA-CoAs are then esterified into phosphatidylethanolamines (PE) and other phospholipids by lysophosphatidylcholine acyltransferase 3 (LPCAT3) or other members of this family ^15^ which, when oxidized and not cleared by GPX4, serve as a death signal for ferroptosis execution ^13, 14^. However, despite these emerging insights into the molecular mechanisms, a better understanding of cellular and metabolic states that confer sensitivity or resistance to ferroptosis is of utmost importance in order to fully exploit its therapeutic potential both in degenerative disease and cancer.

Here, we report that cell density induces a metabolic switch that sensitizes breast cancer cells to ferroptosis. Independently of the cellular phenotype, oncogenic signaling and GPX4/ACSL4 expression, we observe that high cell density confers robust resistance to ferroptosis triggered by pharmacological inhibition or genetic inactivation of GPX4. Mechanistically, we demonstrate that low cell density leads to an enrichment of long-chain and highly unsaturated neutral triacylglycerides (TAG). Furthermore, *de novo* lipogenesis and desaturation pathways were found to be reduced at low cell density, indicating increased fatty acid uptake. The increase in TAG abundance might occur as an adaptive mechanisms to mitigate PUFA-mediated toxicity, which in turn presents a vulnerability towards ferroptosis activation.

## Results

### Cell density sensitizes mammary epithelial cells to ferroptosis irrespective of cellular state

Since recent studies indicate that therapy-resistant cancer cells and those undergoing epithelial-mesenchymal transition (EMT) may become vulnerable to ferroptosis activation ^5–7^, there is growing interest to understand the molecular underpinnings driving ferroptosis sensitivity ^4^. Therefore, we set out in this work to assess whether immortalized human mammary epithelial cells (HMLE) were sensitive to induction of ferroptosis. We also included HMLE-Twist1 cells constitutively expressing the twist family bHLH transcription factor 1 (TWIST1) that, as a consequence, have undergone an epithelial-mesenchymal transition (EMT). EMT programs have been shown to confer resistance to conventional therapies ^16, 17^ and TWIST1 expression has been linked to EMT-activation and metastatic dissemination ^18–20^. As described previously, HMLE cells display an epithelial morphology and express the epithelial marker E-cadherin, whereas *Twist1*-overexpression induces an EMT, resulting in downregulation of E-cadherin, expression of the mesenchymal marker zinc finger E-Box binding homeobox 1 ZEB1 and a mesenchymal morphology (Fig. 1a, d)^18^.

**Fig. 1.**
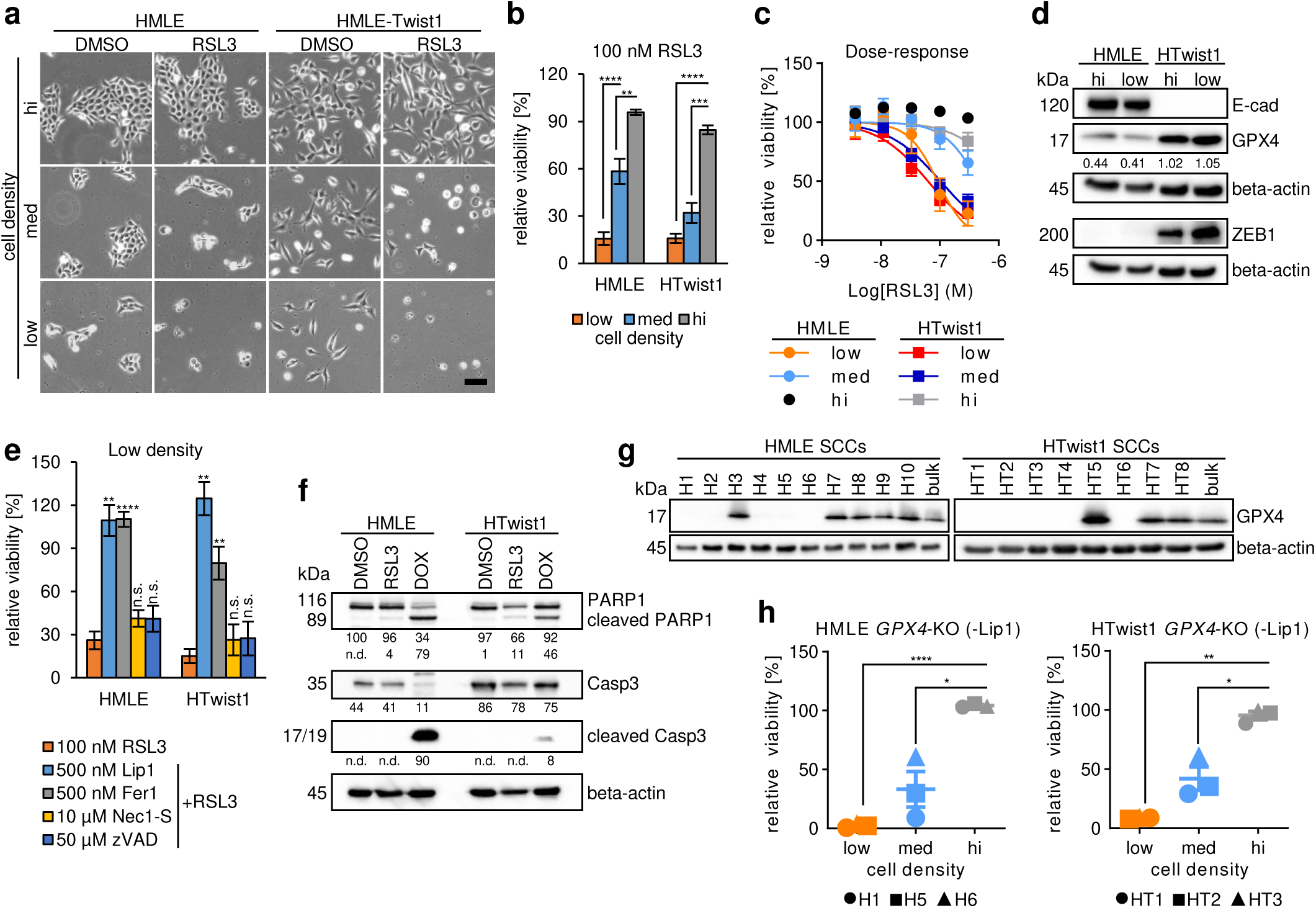
Cell density sensitizes mammary epithelial cells to ferroptosis irrespectively of cellular phenotype. **a** Bright-field microscopy: HMLE and HMLE-Twist1 cells were seeded at three different densities for 24h, then treated with 0.1% DMSO or 100 nM RSL3 for 24h prior imaging. Scale bar: 200 µm. **b** Viability assay: treatment of HMLE and HMLE-Twist1 (HTwist1) with 100 nM RSL3 in a cell density-dependent manner, n=6. **c** Dose-response curves: treatment of HMLE and HMLE-Twist1 seeded at indicated densities with 3-fold dilutions of RSL3, n=4. **d** Immunoblot: E-cadherin (E-cad), GPX4, ZEB1 and beta-actin protein expression in HMLE and HMLE-Twist1 (HTwist1) cells at high (hi) and low density. Upper beta-actin serves as loading control for E-cad and GPX4, lower beta-actin for ZEB1. Numbers indicate densitometric ratios of detected GPX4 protein to beta-actin band in percent. kDa = kilo Dalton. **e** Rescue-viability assay: treatment of HMLE and HMLE-Twist1 cells with DMSO control, 500 nM ferrostatin (Fer1), 500 nM liproxstatin1 (Lip1), 10 µM necrostatin (Nec1-S), 50 µM zVAD-fmk (zVAD) or 100 nM RSL3 alone or in a combination with RSL3 at low density, mean of at least three biological replicates is shown (n=3-5). **f** Immunoblot: protein expression of cleaved and total PARP1, cleaved caspase 3 and caspase 3 (Casp3) and beta-actin in HMLE and HMLE-Twist1 cells upon DMSO, 100 nM RSL3 or 10 µM Doxorubicin (DOX) treatment at intermediate density. Doxorubicin serves as positive control for PARP1 and Casp3 cleavage. beta-actin serves as loading control. From top to bottom, numbers indicate densitometric ratios of detected protein PARP1, cleaved PARP1, Casp3 and cleaved Casp3 to beta-actin band in percent. kDa = kilo Dalton. n.d. = not detectable. **g** Immunoblot: GPX4 and beta-actin protein expression in single cell clones (SCCs) upon CRISPR/Cas9-mediated modification in the *GPX4* locus and parental bulk HMLE and HMLE-Twist1 cells. beta-actin serves as loading control. kDa = kilo Dalton. **h** Viability assay: Lip1 withdrawal of SCCs with *GPX4*-KO seeded at the indicated densities, related to Fig. 1g, n=2 for HMLE and n=4 for HMLE-Twist1 SCCs. Data are presented as mean of indicated biological replicates ± s.e.m. (n=x). Viability was normalized to respective DMSO control (**b**, **c**, **e**) or to respective Lip1 control (**h**) within each seeding density and cell line. Statistics: two-tailed, unpaired T-test with Welch’s correction (p-value: *<0.05, **<0.01, ***0.001, ****<0.0001, n.s. = not significant).

To address whether HMLE and HMLE-Twist1 cells present different susceptibility to ferroptosis, both cell lines were plated at three different cell densities ranging from sparse to sub-confluent and then concomitantly treated with *(1S, 3R)*-RSL3 (RSL3), a GPX4 inhibitor (Fig. 1a, b) ^8^. We found that RSL3 triggered cell death in a cell density-dependent manner with cells seeded at low density being highly sensitive to ferroptosis while those plated at high density being highly resistant (Fig. 1b, c). Moreover, cells plated at an intermediate cell density also showed moderate induction of cell death. We repeated these experiments using cells with a conditional Twist1- or Snail1 (Snail family transcriptional repressor 1)-construct, the latter being another EMT-transcription factor ^18, 21^. Again, we observed density-dependent cell death upon RSL3-treatment in HMLE cells where Twist1 or Snail1 was induced for 15 days, resulting in a full EMT (Supplementary Fig. 1a, b). As a control, *(1R, 3R)*-RSL3, an inactive diastereoisomer of RSL3, did not induce cell death at any tested cell density (Supplementary Fig. 1a). Immunoblotting for GPX4 revealed that HMLE-Twist1 cells expressed 2.3-2.6 fold higher levels of GPX4 protein than HMLE cells, while cell density itself did not have any effect on GPX4 expression levels (Fig. 1d). Together, these data suggest that cell density is a critical factor sensitizing cells to GPX4 inhibition independently of whether HMLE cells were in an epithelial or mesenchymal state, thus contrasting with the recent finding that EMT predisposes cells to ferroptosis ^6^. To investigate whether ferroptosis was the underlying modality of cell death, we treated cells at low seeding density with the pan-caspase inhibitor zVAD-fmk (zVAD) ^22^, or the necroptosis inhibitor Nec1-S that targets the receptor-interacting protein kinase 1 (RIPK1)^23^. We included ferrostatin1 (Fer1) and liproxstatin1 (Lip1) as known inhibitors of ferroptosis ^1, 10^. Indeed, we observed that RSL3-induced cell death was only rescued by treatment with Fer1 or Lip1, but not zVAD or Nec1-S, suggesting that neither apoptosis nor necroptosis are involved (Fig. 1e, Supplementary Fig. 1c). In addition, immunoblotting showed no cleavage of caspase 3 or its downstream target poly(ADP-Ribose) polymerase 1 (PARP1) upon RSL3-treatment in both HMLE and HMLE-Twist1 cells, further ruling out apoptosis as the underlying cell death modality (Fig. 1f) ^24, 25^.

To confirm that ferroptosis-induction at low cell density was exclusively dependent on GPX4 inhibition and not related to off-target effects or simply overall changes in the ratio of inhibitor molecules and cell number, we specifically deleted *GPX4* in HMLE and HMLE-Twist1 cells using CRISPR/Cas9 technology. We derived several single cell clones (SCCs) with and without detectable GPX4 expression (*GPX4*-WT and *GPX4*-KO, respectively, Fig. 1g). Since the genetic knockout of *GPX4* was previously shown to be lethal ^9, 10^, SCCs were kept in Lip1-containing growth medium. In addition, treatment with Fer1 maintained viability as well (Supplementary Fig. 1d, e). Upon withdrawal of Lip1, we again observed induction of cell death only at a low cell density in three *GPX4*-KO SCCs of both HMLE and HMLE-Twist1 cells (Fig. 1h), similar to what was reported for tamoxifen-inducible *Gpx4* knockout fibroblasts ^9^. Importantly, SCCs with intact GPX4 expression still showed cell density-dependent cell death upon RSL3 treatment, suggesting that selected SCCs are representative of the respective bulk population (Supplementary Fig. 1f, g). In conclusion, we observed that cell density determines sensitivity to ferroptosis in both epithelial and *Twist1*-expressing mesenchymal HMLE cells.

### Cell density-dependent cell death is not affected by oncogenic signaling and also occurs in primary mammary epithelial cells

Next, we addressed whether oncogenic signaling affects density-dependent sensitivity to ferroptosis. However, both HMLE cells overexpressing HRas or neuNT oncogenes and HMLE-Twist1 cells overexpressing HRas remained resistant to RSL3-induced cell death at high seeding densities and died at low cell densities, similar to their parental cells (Fig. 2a, Supplementary Fig. 2a). In addition, we tested whether deletion of the tumor suppressor *PTEN* altered sensitivity to RSL3-treatment, but again, we failed to detect differences beyond the cell density effects (Fig. 2b, Supplementary Fig. 2b). In conclusion, neither oncogenic signaling nor an epithelial or a mesenchymal cell state impacted cell density-dependent cell death. Interestingly, investigation of an association between the mutational status of RAS and ferroptosis sensitivity in 117 cancer cell lines did not reveal a selective lethality ^8^, further supporting the notion that still poorly understood and reversible, non-mutational mechanisms impact ferroptosis sensitivity.

**Fig. 2.**
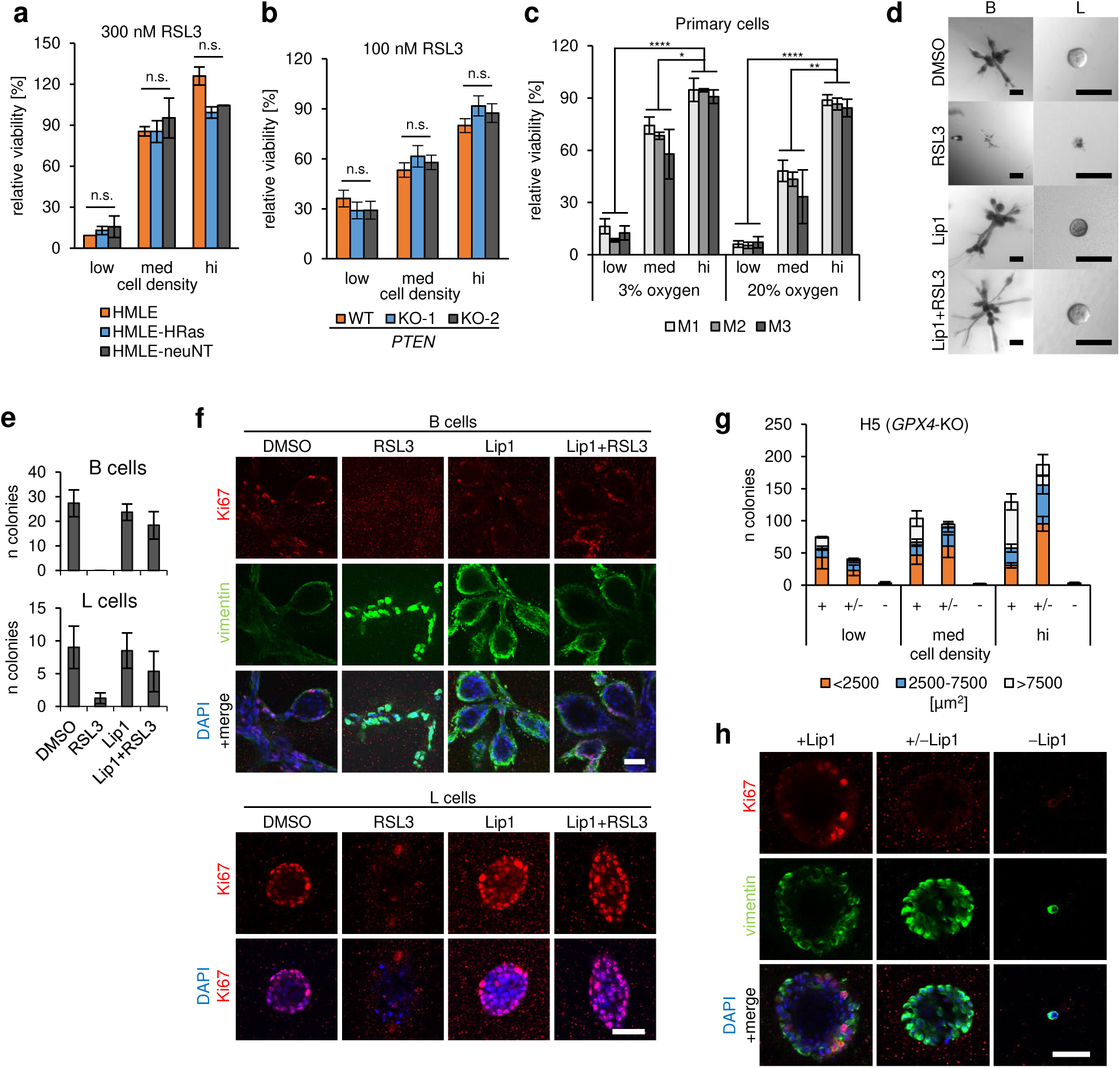
Cell density-dependent cell death is maintained during cellular transformation and prevents growth in 3D-collagen gels. **a** Viability assay: treatment of HMLE, HMLE-HRas (G12V), HMLE-neuNT with 0.3% DMSO or 300 nM RSL3 cells in density, n=2. **b** Viability assay: treatment of *PTEN*-wildtype (WT) and two *PTEN*-knockout clones (KO-1 and KO-2) of HMLE-Twist1-ER cells (without Twist1-activation) in density, n=3. **c** Viability assay: treatment of bulk primary mammary epithelial cells of three different Donors (M1, M2, M3) with 0.1% DMSO or 100 nM RSL3 at ambient oxygen level (20%) or oxygen levels present in tissues (normoxia, 3%) in density, n=3. **d** Bright-field images of representative 3D-collagen gels: sorted CD10-positive basal (B) or luminal progenitor (L) primary mammary epithelial cells were treated with 0.1% DMSO, 100 nM RSL3, 500 nM Lip1 or a combination of RSL3 and Lip1 for 7-10d prior to imaging and quantification of arising colonies. Scale bar: 200 µm. **e** 3D-collagen gels: quantification of colonies as described in Fig. 2d. The mean of 3-4 technical replicates ± s.d. of one representative experiment from two independently performed experiments is shown. **f** Confocal microscopy of 3D-collagen gels: staining of colonies quantified in Fig. 2e with Ki67 (red), vimentin (green) or DAPI (blue, nuclear staining). Scale bar: 50 µm. **g** 3D-collagen gels: HMLE single cell clone H5 with *GPX4*-knockout (KO) was seeded at different densities in 3D collagen gels in medium containing 500 nM Lip1 (+Lip1), withdrawn of Lip1 either directly (-Lip1) or after 3d in initial +Lip1 culture (+/-Lip1) for 7-10 days. Mean of 3-4 technical replicates ± s.d. of one representative experiment from two independently performed experiments is shown. **h** Confocal microscopy of 3D-collagen gels: staining of colonies derived from the intermediate density with Ki67 (red), vimentin (green) or DAPI (blue, nuclear staining). Scale bar: 50 µm. Data (**a-c**) are presented as mean of indicated biological replicates ± s.e.m. (n=x) and were normalized to respective DMSO control within each density and cell line. Statistics: two-tailed, unpaired T-test with Welch’s correction (p-value: *<0.05, **<0.01, ***0.001, ****<0.0001, n.s. = not significant).

To determine whether cell density-dependent ferroptosis sensitivity constitutes an intrinsic property of mammary epithelial cells, primary human mammary epithelial cells (HMECs) isolated from three different donors (M1, M2 and M3) were treated with RSL3 (Fig. 2c). In addition, we included prospectively isolated primary cells of the basal lineage ^26^ to dissect whether lineage identity had an effect on sensitivity to ferroptosis induction (Supplementary Fig. 2c). Again, we observed cell density-dependent cell death induced by RSL3 to a similar extent in all samples. Importantly, this effect occurred both in ambient oxygen atmosphere (20%) as well as 3% oxygen, the latter mimicking physiological tissue pressure (Fig. 2c, Supplementary Fig. 2c). Together, these data indicate that cell density-dependent sensitivity to ferroptosis is a trait present in primary HMECs and does not arise as an artefact of long-term cell culture and selection.

### GPX4 inhibition prevents organoid generation in 3D-collagen gels

Next, we wished to determine whether density-dependent sensitivity to ferroptosis could be observed when cells are grown in a 3D environment. For this purpose, we plated primary HMEC into an organoid assay as previously described ^26^. Specifically, bulk primary HMEC as well as prospectively isolated cells of the basal (B, CD10^+^/CD49f^hi^/EpCAM^−^) and luminal progenitor lineage (L, CD10^−^/CD49f^+^/EpCAM^+^) were plated into 3D-collagen gels. After initial establishment of cultures from single cells and before the onset of organoid formation, RSL3, Lip1 or a combination of both were added to the growth medium every 2-3 days. In the DMSO control and Lip1-treated cells, basal cells gave rise to branched organoids and luminal cells formed spheres, while bulk cells gave rise to different kind of colonies including branched organoids and spheres (Fig. 2d, e, Supplementary Fig. 2d), as reported previously ^26^. RSL3 treatment strongly inhibited colony formation, an effect that was partially rescued by concomitant treatment with Lip1 (60-100% of control, Fig. 2e, Supplementary Fig. 2d). In addition, in contrast to control and rescued samples, we observed a complete absence of proliferation marker Ki67 in the few remaining small clusters of cells in RSL3-treated cultures (Fig. 2f). Together, these data suggest that primary HMEC are highly sensitive to GPX4 inhibition during organoid formation from single cells.

To provide further support for these observations, HMLE *GPX4*-KO SCCs were seeded at different densities in 3D-collagen gels in medium with (+Lip1) or without Lip1 (−Lip1) or cultured for three days in Lip1-containing medium with subsequent Lip1 withdrawal (+/−Lip1, Fig. 2g). This allowed us to test whether cells remain sensitive to ferroptosis once small colonies have formed. As previously described in HMLE cells, *GPX4*-KO SCCs generated multicellular spheres in Lip1-containing medium (Supplementary Fig. 2e) ^21^. Similar to low seeding densities in 2D cultures, immediate withdrawal of Lip1 in 3D cultures resulted in complete growth inhibition (Fig. 2g, Supplementary Fig. 2e). In contrast, Lip1 withdrawal after three days of 3D culture enabled colony formation in similar numbers compared to culture in Lip1-containing growth medium. These data indicate that once colonies have formed, they acquire resistance to cell death induction by *GPX4* knockout. However, both colony size as well as proliferation assessed by Ki67 staining were reduced under this condition, suggesting that Lip1 withdrawal at later time points restricts further expansion in 3D cultures (Fig. 2g, h). Together, these data indicate that, similar to low seeding densities in 2D culture, both primary HMEC as well as HMLE cells are sensitive to ferroptosis at low cell density in 3D culture. However, the observation that HMLE *GPX4*-KO SCCs were partially protected from ferroptosis upon Lip1 withdrawal once small colonies had formed, suggests an intrinsic mechanism that protects epithelial tissues from ferroptosis. In line with these data, a recent study showed that in immortalized non-tumorigenic MCF10A cells and a panel of breast cancer cell lines, detachment of cells from the extracellular matrix (ECM) triggers ferroptosis ^27^. Further, it has been shown that tamoxifen-inducible, c-myc/Hras-transformed *GPX4*-knockout fibroblasts are capable of forming tumors when implanted subcutaneously in mice at very high numbers (5 million)^28^.

### Cell density-dependent ferroptosis does not show all classical hallmarks of ferroptosis and is independent of ACSL4

Ferroptosis has been linked to the degree of unsaturation of PUFAs in cellular membranes, which are particularly prone to peroxidation (Fig. 3a) ^13, 14, 29^. To determine whether levels of oxidized lipids correlated with ferroptosis sensitivity, we measured lipid-derived reactive oxygen species (ROS) levels using the fluorescent dye BODIPY 581/591 C11 ^30^. Consistent with previous observations ^1, 8, 10^, we saw an increase in BODIPY 581/591 C11 fluorescence in *GPX4* knockout cells upon Lip1-withdrawal in both HMLE and HMLE-Twist1 cells at all densities (Fig. 3b, c, Supplementary Fig. 3a). However, this increase in BODIPY 581/591 C11 oxidation greatly varied between different SCCs (Fig. 3b), although all of them were similarly sensitized to ferroptosis at low seeding densities (Fig. 1h). On average, BODIPY 581/591 C11 fluorescence increased at least 1.5-fold upon Lip1-withdrawal in HMLE *GPX4*-KO SCCs irrespective of seeding density (Fig. 3c). Interestingly, HMLE-Twist1 *GPX4*-KO SCCs seeded at an intermediate density showed the strongest accumulation of BODIPY 581/591 C11 fluorescence (4.6-fold on average, Fig. 3c). In summary, Lip1 withdrawal led to an increase in lipid-derived ROS levels in HMLE and HMLE-Twist1 *GPX4*-KO SCCs in all seeding densities. Therefore, the observed increase in BODIPY 581/591 C11 oxidation may not necessarily correlate with the induction of ferroptosis. Consequently, these data suggest that cells at high and low seeding densities may cope differently with an increase in lipid-derived ROS levels, rather than cell density having a strong impact on overall lipid peroxidation levels. To further characterize cell density-dependent ferroptosis, we undertook a series of experiments modulating PUFA-incorporation and -peroxidation. Recent studies showed that ACSL4, which plays a key role in lipid biosynthesis and fatty acid degradation, is an essential enzyme for ferroptosis execution ^13, 14^. ACSL4 increases the PUFA-content within phospholipids, which are susceptible to oxidation. Consequently, blocking ACSL4 activity by rosiglitazone (ROSI) has been shown to protect against ferroptosis ^13^. Lip1, Fer1, or the lipophilic antioxidant α-tocopherol (α-toc) can also prevent accumulation of lipid oxygenation and thus, execution of ferroptosis. Moreover, since the labile iron pool (redox-active Fe^2+^) contributes to lipid oxygenation via the Fenton reaction, chelation of iron by deferoxamine (DFO) or cyclopirox (CPX) has been shown to protect against ferroptosis (Fig. 3a) ^1^.

**Fig. 3.**
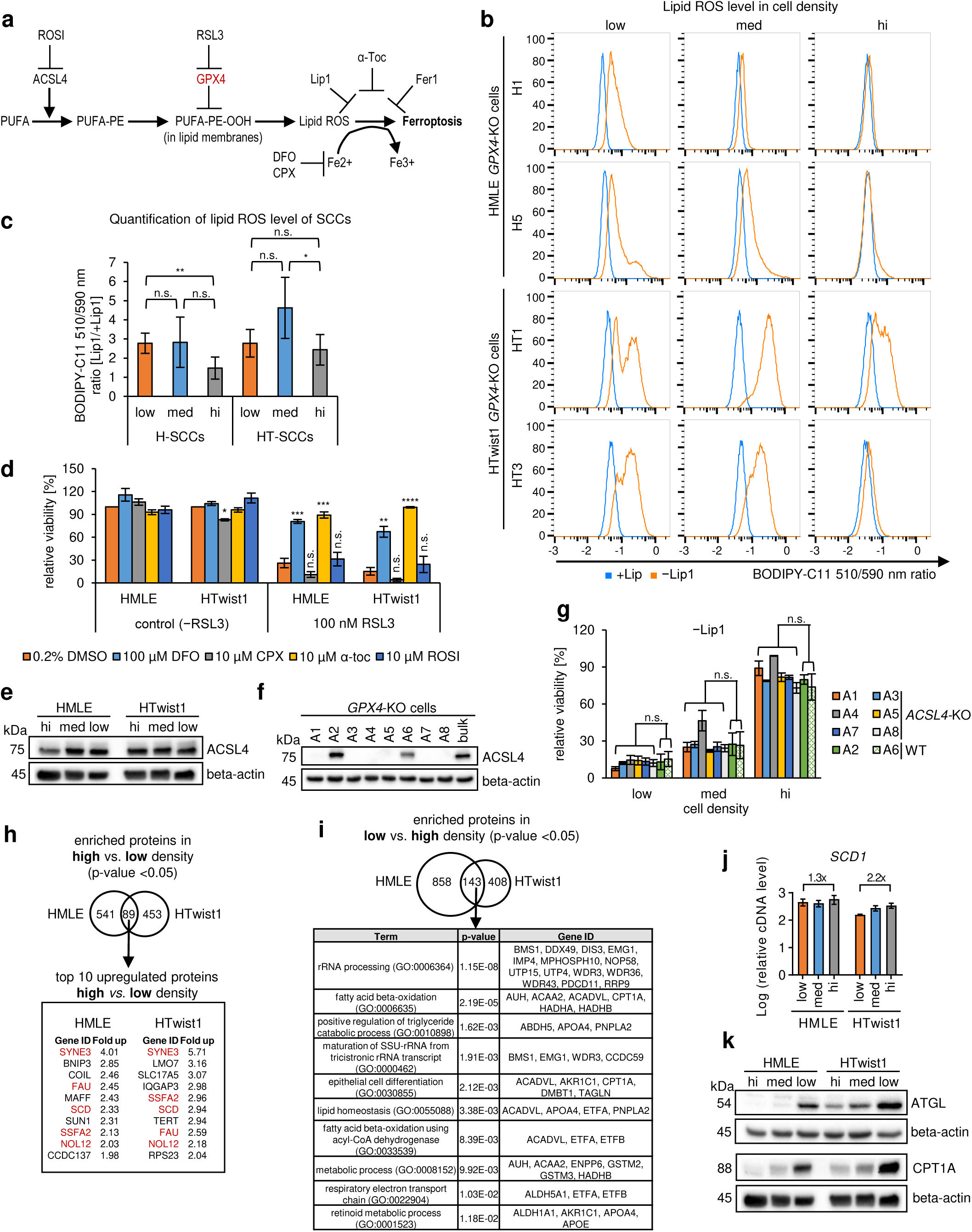
Cell density-dependent ferroptosis does not show all classical hallmarks of ferroptosis and is independent of ACSL4. **a** Scheme showing regulators of ferroptosis and their inhibitors, GPX4: glutathione peroxidase 4, ACSL4: acyl-CoA synthetase long chain family member 4, ROSI: Rosiglitazone, PUFA: polyunsaturated fatty acid, PE: phosphatidylethanolamine, -OOH: hydroperoxide, α-toc: α-tocopherol, DFO: deferoxamine, CPX: ciclopirox, Lip1: liproxstatin1, Fer1: ferrostatin1, ROS: reactive oxygen species. **b** Flow cytometry: BODIPY-C11 staining of *GPX4*-knockout SCCs seeded at different densities in 1 µM Lip1 medium (+Lip1, blue) or in medium without Lip1 (−Lip1, orange). X-axis: log10 of BODIPY-C11 510/590 nm fluorescence ratio, Y-axis: percentage of the maximum count, one representative experiment showing variation in fluorescence level is shown. **c** Quantification of BODIPY-C11 510/590 nm fluorescence ratio of HMLE (H) and HMLE-Twist1 (HT) single cell clones (2 clones of H-SCCs and 3 clones of HT-SCCs, respectively) upon *GPX4*-knockout (−Lip1) normalized to respective +Lip1 control seeded and stained as described in Fig. 3b. n=6. **d** Rescue-viability assay: treatment HMLE and HMLE-Twist1 cells with the indicated compounds as described in Fig. 3a alone or in a combination with 100 nM RSL3 at low density, mean of at least three biological replicates is shown (n=3-5). **e** Immunoblot: ACSL4 and beta-actin (loading control) protein expression in HMLE and HMLE-Twist1 cells seeded in density, kDa = kilo Dalton. **f** Immunoblot: ACSL4 and beta-actin (loading control) protein expression in single cell clones (SCCs) of HMLE H5 *GPX4*-KO cells derived upon CRISPR/Cas9-mediated modification in the *ACSL4* locus. kDa = kilo Dalton. **g** Viability assay: deprivation of 1 µM Lip1 in CRISPR/Cas9-derived SCCs with *ACSL4*-knockout (*ACSL4*-KO) or with intact *ACSL4*-expression (*ACSL4*-WT) in density, n=3. **h** Venn diagram of proteomic data sets representing significantly, upregulated proteins (p-value: <0.05) at high density compared to low density in HMLE (left, 630 proteins) and HMLE-Twist1 (right, 542 proteins) cells. The overlapping 89 proteins are significantly, upregulated under high cell density conditions in both cell lines. Top 10, significantly enriched proteins at high cell density compared to low cell density for each cell line of the commonly, upregulated data set (89 proteins) are shown. n=4. **i** Venn diagram of proteomic data sets representing upregulated proteins (p-value: <0.05) at low density compared to high density in HMLE (left, 1001 proteins) and HMLE-Twist1 (right, 551 proteins) cells. The overlapping 143 proteins are significantly upregulated under low cell density conditions in both cell lines. GO-term analyses: top 10, significantly enriched terms (biological processes) of common upregulated proteins (143) in low cell density. n=4. **j** RT-qPCR: *SCD1* mRNA expression of HMLE and HMLE-Twist1 cells seeded in density. *RPL32* was used as an internal control. Mean of three technical replicates ± s.e.m. of one representative experiment from two independently performed experiments is shown. Y-axis: log10. **k** Immunoblot: ATGL, CPT1A and beta-actin (loading control) protein expression in HMLE and HMLE-Twist1 cells seeded in density. kDa= kilo Dalton. Data are presented as mean of indicated biological replicates ± s.e.m. (n=x). Data was normalized to respective DMSO control (**d**) or to respective +Lip1 control (**c, g**) within each seeding density and cell line. Statistics: two-tailed, unpaired T-test with Welch’s correction (p-value: *<0.05, **<0.01, ***0.001, ****<0.0001, n.s. = not significant).

To understand why cells die in a cell density-dependent manner, we tested the ability of the aforementioned inhibitors to rescue RSL3-induced cell death. We observed that iron-chelation by DFO, but not CPX, partially rescued RSL3-induced cell death in both HMLE and HMLE-Twist1 cells (28% to 72% viability in HMLE and 17% to 55% viability in HMLE-Twist1 cells upon DFO-co-treatment, Fig. 3d). In contrast to DFO, CPX directly chelates iron intracellularly which might affect iron-containing enzymes as well ^31^, suggesting that excessive iron chelation might produce opposing effects. Moreover, the lipophilic antioxidant α-toc rescued RSL3-induced cell death (Fig. 3d). These results thus suggest that peroxidation of PUFA-containing lipids contributes to cell density-dependent cell death (Fig. 3d). To assess whether ACSL4 contributes to the intracellular PUFA-containing lipid pool, cells were pre-treated with ROSI prior to treatment with RSL3. Surprisingly, ROSI treatment did not rescue cell death in HMLE and HMLE-Twist1 cells (Fig. 3d), although both HMLE and HMLE-Twist1 cells expressed ACSL4 at the RNA and protein level at all densities (Fig. 3e, Supplementary Fig. 3b). ROSI is an agonist of the peroxisome proliferation-activated receptor-γ (PPAR-γ), and ACSL4 inhibition is only an off-target effect ^32^. Therefore, to validate whether ACSL4 might be involved in cell density-dependent cell death, we performed a CRISPR/Cas9-mediated knockout of *ACSL4* in HMLE *GPX4*-KO SCCs (Fig. 3f, 1g, h). We derived several clones with *ACSL4* knockout (*ACSL4*-KO: A1, A3, A4, A5, A7 and A8) and control clones with intact ACSL4 expression (A2 and A6 respectively, Fig. 3f). Unexpectedly, all clones died irrespective of ACSL4 expression in a cell density-dependent manner upon withdrawal of Lip1 (Fig. 3g). Similar results were obtained when ACSL4 was deleted in both HMLE and HMLE-Twist1 cells using siRNAs and treated with RSL3 (Supplementary Fig. 3d, e), suggesting that cell density-dependent ferroptosis occurred independently of ACSL4 in both cell lines. Since other ACSL enzymes have been linked to ferroptosis ^33^, knockdown experiments using siRNAs against ACSL1 and ACSL3 were performed. Similar expression levels were observed at different densities for ACSL1 and ACSL3 in HMLE and HMLE-Twist1 cells (Supplementary Fig. 3b, c). However, neither knockdown of ACSL1 nor knockdown of ACSL3 impacted cell density-dependent cell death induced by RSL3 in both HMLE and HMLE-Twist1 cells (Supplementary Fig. 3d, e), indicating that other lipid-related proteins are important for cell density-dependent ferroptosis. Together, cell density-dependent ferroptosis shows some hallmarks of ferroptosis such as iron dependency and lipid peroxidation, but we could not establish a correlation of cell death induction and upregulation of global lipid peroxidation nor a dependence of cell death on ACSL4.

### Cell density regulates lipid metabolism-related proteins

To further explore the mechanisms of the cell density-dependent effects toward ferroptosis, we performed a proteomic study with four independent biological replicates to assess which proteins were differentially expressed at high and low cell densities with the rationale that these proteins might be potentially involved in cell density-dependent cell death. For data analysis, we filtered for significantly enriched proteins at high compared to low cell density in HMLE cells (630 proteins, p<0.05) or in HMLE-Twist1 cells (542 proteins, p<0.05) and then, we assessed the upregulated overlapping proteins at high cell densities (89 proteins) between the two cell lines (Fig. 3h, Supplementary Table 1). Moreover, we performed the same analysis to filter for enriched proteins at low compared to high cell density for HMLE cells (1001 proteins, p<0.05) and HMLE-Twist1 cells (551 proteins, p<0.05) prior to selection of the upregulated proteins at low cell densities (143 proteins) shared by both cell lines (Fig. 3i, Supplementary Table 1). Notably, the fold regulation and thus most regulated proteins partly differed between HMLE and HMLE-Twist1 cells, although some proteins as for example the stearoyl-CoA desaturase-1 (SCD1) or apolipoprotein E (APOE) were found within the top ten regulated proteins in both cell lines (Fig. 3h, Supplementary Fig. 3f). The list of enriched proteins at low or high cell density in both cell lines was submitted to GO-term enrichment analysis (biological processes) using DAVID (Fig. 3i, Supplementary Fig. 3g, Supplementary Table 1) ^34, 35^. Interestingly, several GO-terms related to lipid metabolism were enriched at both cell densities (Fig. 3i, Supplementary Fig. 3g). For example, the GO-term “long-chain fatty-acyl-CoA biosynthetic process” was enriched at high cell density (Supplementary Fig. 3g) and by RT-qPCR we could verify a slight upregulation of SCD1 at high cell density in both cell lines (Fig. 3j), while ACSL3 was only upregulated at high cell density in HMLE-Twist1 cells (Supplementary Fig. 3b). SCD1 is an enzyme required for monounsaturated fatty acid (MUFA) synthesis by catalyzing the desaturation of saturated fatty acids ^36^. At low cell density, GO-terms like “fatty acid beta-oxidation”, “positive regulation of triglyceride catabolic process”, “lipid homeostasis” and “respiratory electron transport chain” were enriched (Fig. 3i). To confirm cell density-dependent regulation of selected proteins, we validated adipose triglyceride lipase (ATGL = *PNPLA2*), which is the initial enzyme for triacylglyceride (TAG) hydrolysis and is regulated by its co-activator ABHD5 ^37^. Functionally, ATGL catalyzes TAGs from intracellular lipid droplets (LDs), reducing LD abundance ^38^. We validated ATGL expression by immunoblotting and observed an upregulation of ATGL in both HMLE and HMLE-Twist1 cells at low compared to high seeding densities (Fig. 3k). Furthermore, we observed that enzymes required for β-oxidation such as carnitine palmitoyltransferase 1A (CPT1A) or acetyl-CoA acyltransferase 2 (ACAA2) were upregulated at low compared to high density in each line (Fig. 3k, Supplementary Fig. 3h). Together, these data indicate that lipid metabolism is regulated by cell density and further suggest that low cell density might upregulate TAG metabolism and oxidative metabolism.

### Epithelial and Twist1-induced mesenchymal HMLE cells enrich polyunsaturated triacylglycerides at low cell density

Since the proteomic data indicated that changes in lipid metabolizing genes might confer sensitivity to ferroptosis at low density in breast cancer cells, we analyzed the lipidome of cells seeded at low, intermediate and high cell density. Principal component analysis (PCA), calculated based on all the measured lipid species across all lipid classes, revealed distinct lipid profiles at different cell densities (Fig. 4a). Lipid profiles were most distinct between high and low cell density while lipid profiles of epithelial HMLE and Twist1-induced mesenchymal cells seeded clustered within each cell density. Interestingly, the lipid profile of HMLE cells seeded at an intermediate cell density clustered closer to high cell density lipid profiles of both cell lines, while HMLE-Twist1 cells seeded at an intermediate cell density showed a lipid profile in between of cells seeded at high or low cell density (Fig. 4a). Furthermore, clustering of lipid profiles by PCA correlates to ferroptosis-sensitivity of HMLE and HMLE-Twist1 cells (Fig. 1b, c). Both epithelial and mesenchymal cells are highly resistant to ferroptosis at high cell density while sensitive at low cell density. At intermediate cell densities, HMLE-Twist1 cells are more sensitive than HMLE cells, suggesting that changes in lipid metabolism and thus lipid profiles confer resistance or sensitivity to ferroptosis. However, cell density did not influence the abundance of most detected lipid classes nor of phospholipids like PEs that has been associated with ferroptosis ^13, 14^ (Fig. 4b). In contrast, we observed a significant increase in TAGs at low cell density in both cell lines (Fig. 4c), which constitute the major components of LDs. Immunofluorescent staining of TAGs using LipidTOX Green indicated a localization of TAGs in LDs at low cell density in contrast to high cell density where the staining appeared to be diffusely located around the nucleus (Supplementary Fig. 4a). Furthermore, at low cell density, TAGs were significantly enriched with long-chain fatty acids that showed a higher degree of unsaturation (Fig. 4d, e, Supplementary Fig. 4b-d). For example, PUFAs like arachidonic acid (AA, C20:4) or adrenic acid (AdA, C22:4) were elevated in TAGs at low cell density (Supplementary Fig. 4d). In contrast, TAGs esterified with saturated fatty acids (SFA) like palmitate (C16:0) or monounsaturated fatty acids (MUFAs) like oleate (C18:1) were overrepresented in TAGs at high cell density (Fig. 4d, e, Supplementary Fig. 4a-c). Interestingly, we observed a similar trend of PUFA-enrichment in phospholipids like PEs, phospatidylcholines (PCs), phosphatidylglycerols (PGs) and phosphatidylinositols (PIs) at low cell density and an enrichment of SFA/MUFAs-containing phospholipids at high cell density (Fig. 4f, Supplementary Fig. 4e-g). In line, peroxidation of AA and AdA-containing PEs were identified as death signals in ferroptosis execution ^13, 14^, yet the striking increase in TAG abundance and in particular PUFA-TAG abundance points to a central role of TAGs in cell density-dependent ferroptosis sensitivity. Moreover, the data suggest that alterations in fatty acid metabolism and saturation by cell density, i.e. SFA/MUFA and PUFA metabolism, may be a driver of cell density-dependent ferroptosis.

**Fig. 4.**
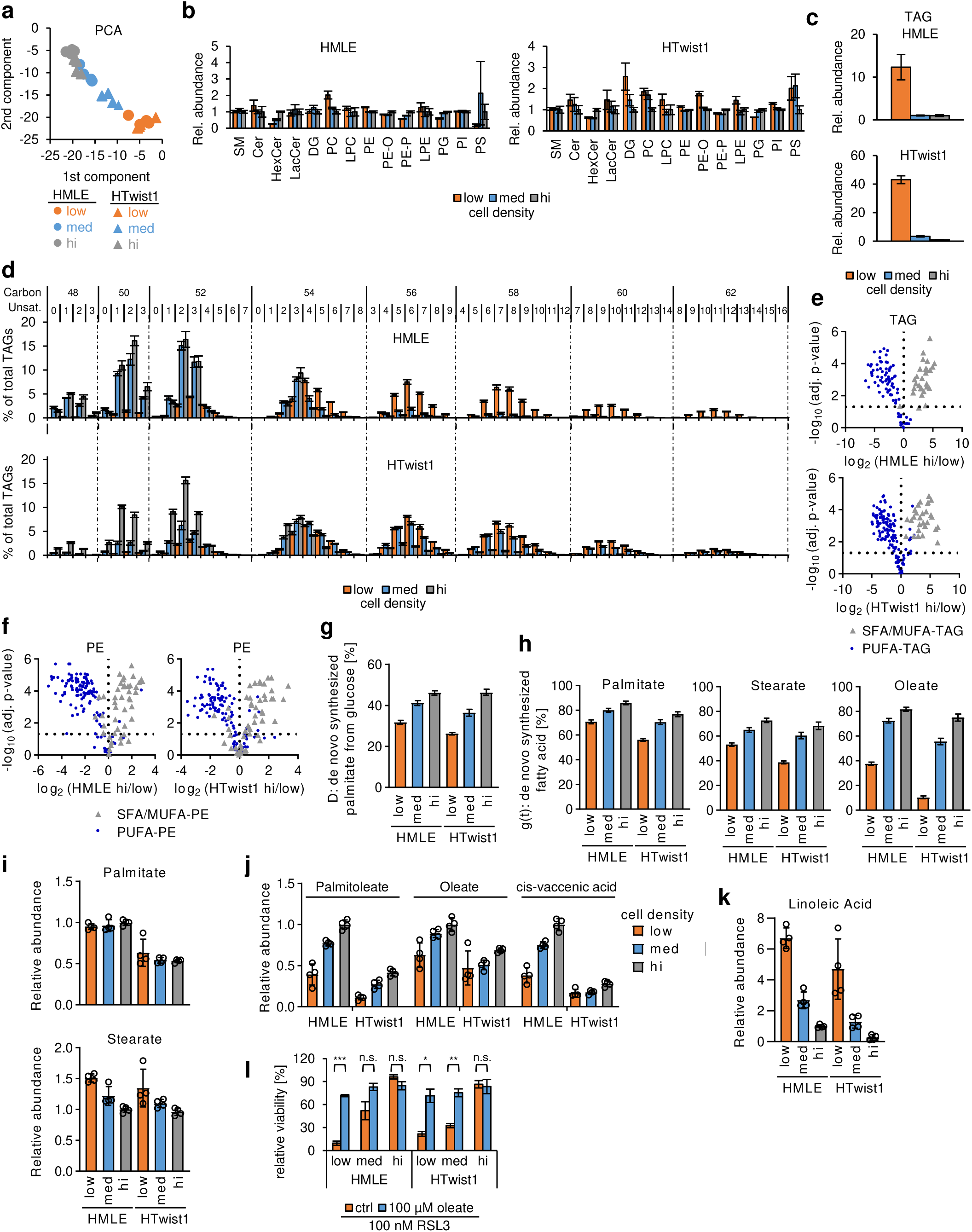
Epithelial and Twist1-induced mesenchymal HMLE cells enrich polyunsaturated triacylglycerides at low cell density. **a** Principal component analysis (PCA) of lipid profiles of HMLE and HMLE-Twist1 cells seeded at density. n=4. **b** Relative abundance of indicated lipid classes of HMLE and HMLE-Twist1 cells seeded at density. SM: sphingomyelins, Cer: ceramides, HexCer: hexosylceramides, LacCer: lactosylceramides, DG: diacylglycerides, PC: phosphatidylcholine, LPC: lysophosphatidylcholine, PE: phosphatidylethanolamine, PE-O: 1-alkyl,2-acylphosphatidyethanolamines, PE-P: 1-alkenyl,2-acylphosphatidyethanolamines, LPE: lysophosphatidylethanolamine, PG: phosphatidylglycerol, PI: phosphatidylinositol, PS: phosphatidylserine. n=4. **c** Relative abundance of triacylglycerides (TAG) of HMLE and HMLE-Twist1 cells seeded at density. n=4. **d** Relative TAG abundance at density of HMLE and HMLE-Twist1 cells denoted by sum notation and sorted from left to right by carbon chain length and by saturation degree within groups of identical chain length. n=4. **e** Volcano plot: log2 transformed fold changes of relative TAG species abundance between high (hi) and low cell density for each cell line plotted against -log10 FDR adjusted p-values calculated using the Benjamini/Hochberg procedure. TAG species were divided into SFA/MUFA-TAGs (grey triangle) or PUFA-TAGs (blue dot). n=4. **f** Volcano plot: fold changes of relative PE species abundance grouped as SFA/MUFA-PEs (grey triangle) or PUFA-PEs (blue dots) between high (hi) and low cell density for each cell line, plotted as described in Fig. 4e. n=4. **g** Isotopomer spectral analysis (ISA) of ^13^C_6_ glucose contribution to newly synthesized palmitate (D value) shown as percentage. HMLE and HMLE-Twist1 were seeded at density and incubated for 48h with media containing 8 mM ^13^C_6_ glucose. Data show mean ± c.i., n=4. **h** Isotopomer spectral analysis (ISA) for the fraction of newly synthesized palmitate, stearate and oleate (g(t)) shown as percentage. Related to Fig. 4g n=4. **i** Normalized palmitate and stearate abundance in HMLE and HMLE-Twist1 cells seeded at density. n=4. **j** Normalized metabolic abundance of indicated fatty acids in HMLE and HMLE-Twist1 cells seeded at density. n=4. **k** Normalized linoleic acid abundance in HMLE and HMLE-Twist1 cells seeded at density. n=4. l Viability assay: treatment of HMLE and HMLE-Twist1 seeded at indicated densities with 100 nM RSL3 (+RSL3) only (ctrl, orange) or pre-treated for 24h with 100 µM oelate (blue). Oleate was present during DMSO/RSL3 treatment, n=3. Data are presented as mean of indicated biological replicates ± s.e.m. (n=x, **b-d**, **l**) or ± s.d. (**i-k**). Data were normalized to high cell density condition within each cell line (**b, c**) or to HMLE high cell density condition of the respective fatty acid (**i-k**).

To delineate whether massive TAG accumulation at low cell density occurs via *de novo* lipogenesis, we performed ^13^C_6_-glucose labelling experiments for 48h followed by mass spectrometry analysis of incorporated ^13^C carbons into fatty acids. Thereby, we observed that palmitate produced from ^13^C_6_-glucose, which is an initial fatty acid in the *de novo* lipogenesis ^36^, is reduced at low cell density (Fig. 4g) suggesting that the fraction of glucose contributing to *de novo* lipogenesis is reduced at low cell density in both cell lines. Furthermore, we found that the fraction of newly synthesized palmitate is reduced at low cell density in both cell lines (Fig. 4h, Supplementary Fig. 4h). Notably, the reduction in newly synthesized palmitate cannot be explained by different proliferation rates at different densities (Supplementary Fig. 4i-k). Likewise, *de novo* lipogenesis of the elongation product stearate (C18:0) from palmitate was reduced at low cell density (Fig. 4h). Interestingly, the fraction of newly synthesized oleate (C18:1), which is a desaturation product of stearate, was even more reduced at low cell density (Fig. 4h), suggesting that *de novo* desaturation is particularly impaired at low cell density ^39^. Overall, the abundance of palmitate was similar at different cell densities, albeit approximately 1.5-fold lower for HMLE-Twist1 cells (Fig. 4i), indicating that other carbon sources such as glutamine are used for palmitate synthesis or palmitate is used for other metabolic processes at high cell density. We further found that the saturated fatty acid stearate, an elongation product of palmitate, increased in abundance at low cell density compared to high cell density (Fig. 4i), while the levels of further processed, monounsaturated fatty acids such as palmitoleate, oleate and cis-vaccenic acid were all reduced at lower densities in both cell lines (Fig. 4j). These results indicate that desaturation is impaired at low cell density. Since SCD1 is required for these desaturation products (Fig. 4j) ^39^, these data point to a reduction of SCD1 activity at low cell density which is in line with increased SCD1 abundance at high cell density found by proteomics (Fig. 3h). Together, these data suggest that *de novo* lipogenesis and in particular *de novo* MUFA synthesis is decreased at low cell density which indicates that massive TAG accumulation most likely occurs via fatty acid uptake. This is further supported by the observation that levels of the essential fatty linoleic acid were also elevated at low cell density compared to high cell density in both cell lines (Fig. 4k). These data suggest that low cell density leads to an enrichment of fatty acids and in particular PUFA uptake, while *de novo* lipogenesis and in particular desaturation at low cell density is reduced. A recent report showed that PUFA-TAGs stored in LDs reduce lipotoxicity induced by unsaturated fatty acids ^40^, indicating that the massive increase in TAGs might be a compensatory mechanism to reduce lipid toxicity at low cell density. Furthermore, it has been shown that supplementation with oleate or increased SCD1 activity reduced FA-induced lipotoxicity by promoting TAG accumulation ^41^. Interestingly, oleate treatment completely rescued RSL3-induced cell death induced at low and intermediate seeding densities (Fig. 4l), whereas oleate treatment alone only had a minor impact on cell viability (Supplementary Fig. 4l). Supplementation with oleate has been shown to confer a ferroptosis-resistant cell state by decreasing PUFA-containing phospholipids ^33^. This further supports our hypothesis that PUFA uptake might be highly upregulated at low cell density, thereby impacting the PUFA to SFA/MUFA ratio and thus ferroptosis sensitivity. A recent study linked HIF2α, which increased PUFA-enriched lipids and TAGs, to a ferroptosis-susceptible cell state in cancer cell lines with a clear-cell morphology ^42^. Therefore, we assessed the expression of HIF2α at different densities in HMLE and HMLE-Twist1 by immunoblotting. However, protein expression levels of HIF2α were higher at high density in both cell lines compared to low density (Supplementary Fig. 4m), pointing to another mechanism of PUFA enrichment than recently proposed ^42^. Instead our work highlights that cell density highly influences lipid metabolism by regulating lipid metabolism-related enzymes and that, in turn, induces the enrichment of polyunsaturated lipids and in particular PUFA-TAGs at low cell density.

## Discussion

Together, our findings suggest that the switch in lipid metabolism correlating with cell density plays a critical role in determining sensitivity to ferroptosis in both primary mammary epithelial and breast cancer cells. We further found that cell density-dependent ferroptosis occurs irrespectively of the cellular phenotype in epithelial and Twist1-induced mesenchymal breast cancer cells, contrasting a recent finding showing that therapy-resistant, high mesenchymal cell states sensitize cells to ferroptosis ^6^. These seemingly disparate results suggest that studying different cellular contexts is of utmost importance to further shed light on mechanisms that determine ferroptosis sensitivity. For instance, earlier studies provided a link between RAS expression and ferroptosis sensitivity ^43^, which was not necessarily predictive across a large panel of cancer cell lines ^8^, again pointing to different mechanisms of ferroptosis sensitivity in different cellular contexts. Another mechanism of the impact of cell density on ferroptosis reported recently involves E-cadherin-mediated ferroptosis suppression via NF2-YAP signaling ^44^. However, the proposed signaling modulating ferroptosis sensitivity may not apply to our cellular systems since E-cadherin levels in HMLE cells were not affected by cell density and HMLE-Twist1 cells do not express E-cadherin, but still show density-dependent ferroptosis-sensitivity. Instead, both cellular cell states massively accumulated TAGs enriched with PUFA at low cell density. Yet the biological meaning of PUFA-TAG enrichment has to be clarified in future studies. Accumulation of PUFAs in TAGs might serve as sink to reduce ROS and lipid peroxidation as recently proposed ^40^. Interestingly, proliferation of neuronal stem cells during oxidative stress has been shown to involve LD biogenesis in adjacent nice glia cells that sequestered PUFAs to limit ROS and lipid peroxidation ^45^. Along these lines, another recent report described a metabolic coupling of neuron-astrocytes engaging LD formation and oxidative metabolism, thereby protecting neurons from fatty acid toxicity ^46^. Furthermore, metastatic cells, at least in melanoma, have been shown to display increased oxidative stress which has to be limited at the distant site to enable metastatic outgrowth ^47^.

In conclusion, our studies suggest that PUFA-TAG accumulation might be a novel marker for ferroptosis-sensitive cells, which could be mechanistically exploited to target breast cancer cells during stages of metastasis, where they are present as single or small clusters of cells, for example following systemic dissemination into distant tissues.

## Materials and Methods

### Chemicals

The following chemicals were purchased from Sigma: α-tocopherol, Bovine serum albumin (BSA), Ciclopirox, Deferoxamine, Dimethyl sulfoxide (DMSO), Doxorubicine hydrochloride, Ferrostatin-1, Liproxstatin-1, Necrostatin-1S, Oleic Acid-Albumin from bovine serum, Rosiglitazone. RSL3 was purchased from Cayman Chemical. z-VAD-fmk was purchased from R&D Systems.

### Cell culture

Primary mammary epithelial cells (HMECs) were isolated from reduction mammoplasties according to a published protocol ^48^ with modifications as previously described ^26^ in accordance with the regulations of the ethics committee of the Ludwig-Maximilians-University Munich (ethics vote EK397-12). HMLE, HMLE-Twist1, HMLE-Ras, HMLE-Twist1-Ras and HMLE-neu-NT cells were kindly gifted by Robert A. Weinberg (Whitehead Institute). Briefly, immortalized HMLE cells were generated by retroviral transduction of SV40 large T early region and catalytic subunit of human telomerase enzyme (hTERT) ^49^. HMLE-Twist1 were subsequently generated by transduction with a pBabe-Puro-Twist1 vector, leading to constitutive Twist1-overexpression and a mesenchymal phenotype ^19^. HMLE-Ras cells were generated by introduction of a pBabe-Puro-Ras (V12H) retroviral vector ^49^ and HMLE-neu-NT cells with a pWZL-vector containing a mutated form of the HER2/neu oncogene ^18^. Cells were grown in a 37°C incubator with a humidified atmosphere of 5% CO2, except for primary cells, which were maintained at 3% oxygen level if not otherwise stated. All cells were cultured in mammary epithelial cell growth medium containing 0.004 ml/ml BPE, 10 ng/ml EGF, 5 µg/ml Insulin and 0.5 µg/ml hydrocortisone (PromoCell) supplemented with 1% Pen/Strep (Invitrogen) (MECGM). For *GPX4*-knockout single cell clones of HMLE and HMLE-Twists1 cells (for generation see CRISPR/Cas9-mediated knockout of individual genes section), 500 nM to 1 µM Lip1 was additionally added to MECGM. Inducible HMLE-Twist1-ER cells and HMLE-Snail1-ER were kindly gifted by Robert A. Weinberg (Whitehead Institute) ^18^ and cultured in presence of 10 µg/ml blasticidin (Life Technologies). For induction of EMT, cells were additionally treated with 20 nM 4-hydroxy-tamoxifen (TAM, Sigma) for the indicated number of days. To visualize cell death induced by RSL3, HMLE and HMLE-Twist1 cells were plated in 6-well plates at a density of 90000 (high), 30000 (med) and 10000 (low) cells, corresponding to 3000 (high), 1000 (med) and 333 (low) seeded cells per well of 96-wells plates. The next day, cells were treated with 0.1% DMSO or 100 nM RSL3 and 20-24h later, bright-field images were taken on a Leica DM IL LED microscope.

### Viability Assays

To measure the viability of cells in different densities, cells were seeded in 96-wells plates at a density of 3000, 1000 and 333 cells per well (high, med and low respectively, 3-6 technical replicates). Treatment with DMSO control or RSL3 was started one day after plating and cells were treated for 20-24h. For rescue-experiments, cells were seeded at a density of 600 cells per well. Compounds employed in viability assays were added during RSL3 treatment, except for Rosiglitazone and oleate (additionally 24h pre-treatment). For assessment of the viability of CRISPR/Cas9 *GPX4*-knockout clones and *GPX4*/*ACSL4*-knockout clones, Lip1 was withdrawn from the medium directly on the day of plating. For all experiments, viability was measured 48h after plating using CellTiter-Glo assay (Promega). CellTiter-Glo assay is a luciferase-based assay that measures endogenous ATP levels that correlate with the number of metabolically active, and thus viable cells. Luminescence was detected using the Luminometer Centro XS^3^ LB 960 (Berthold Technologies). Obtained RLU values were averaged and normalized to respective DMSO control or Lip1 control.

### Cell proliferation

To measure cell proliferation, HMLE and HMLE-Twist1 cells were plated at 3000, 1000 and 333 cells per well (high, med and low respectively). Proliferation was monitored every 24h over a period of 96h using CellTiter-Glo assay (Promega). Growth medium was refreshed every 24h. Data obtained was normalized to the 24h time point for every cell density within each cell line.

### Cell cycle analysis

For cell cycle analysis, HMLE and HMLE-Twist1 cells were seeded at 516000, 172000 and 57000 cells (high, med and low) in 10-cm dishes. After 48h, cells were harvested by trypsinization, resuspended in PBS and fixed by addition of 100% ethanol. After overnight incubation at -20°C, fixed cells were washed twice with PBS and RNA was removed by RNase A (Thermo Fisher Scientifc) treatment. DNA was stained with propidium iodide (Thermo Fisher Scientific) overnight at 4°C, and after washing with PBS resuspended in FACS buffer followed by analysis on a FACS Aria IIIu (BD Biosciences) using laser and filters according to manufacturer’s instructions. Data was analyzed using FlowJo Software (FlowJo, LLC). A linear scale was used to visualize PI staining, thus cell cycle distribution.

### Immunoblotting

For blotting against GPX4, 1.5 x 10^6^ and 0.3 x 10^6^ cells (high and low) cells were seeded on 15-cm dishes corresponding to 3000 and 600 cells per wells of 96-wells plates. For ATGL, ACSL1, ACSL4, CPT1A and HIF2α protein detection, 516000, 172000 and 57000 cells (high, med and low) were seeded in 10-cm dishes, corresponding to 3000, 1000 and 333 cells in 96-wells. For detection of apoptosis markers, cells were seeded at a density of 500000 cells in 15-cm dishes corresponding to 1000 cells (med) in 96-wells, treated the next day for 20h with 0.1% DMSO, 100 nM RSL3 or 10 µM Doxorubicin. For pAKT and total AKT assessment, seeding density in 10-cm dishes corresponded to 3000 cells in 96-well and for starvation of cells, medium was changed on the second day to basal DMEM/F12 overnight. For all experiments, protein isolation was performed 48h after plating. Briefly, dishes were washed with PBS, cells scraped off on ice, pelleted and resuspended in RIPA buffer (50 mM Tris pH 8.0, 150 mM NaCl, 1% NP-40, 0.5% sodium deoxycholate, 0.1% sodium dodecyl sulfate and 5 mM EDTA pH 8.0) containing phosphatase and protease inhibitor cocktails (Sigma). Protein concentration was measured using the DC Protein Assay (Bio-Rad Laboratories). Protein lysates (10-20 µg) were separated on 12.5% SDS-PAGE gels followed by wet-blot transfer to PVDF membranes. Western blotting was performed using the following primary antibodies: GPX4 (EPNCIR144, ab125066, Abcam, 1:2000), ACSL4 (A5, sc-271800, Santa Cruz, 1:100), E-cadherin (EP700Y, GTX61329, Biozol, 1:25000), Zeb1 (HPA027524, Sigma, 1:5000), phospho-Akt Ser473 (D9E, 4060, Cell Signaling Technology, 1:2000), pan Akt (C67E7, 4691, Cell Signaling Technology, 1:1000), Caspase 3 (9662, Cell Signaling Technology, 1:1000), cleaved Caspase 3 Asp175 (5A1E, 9664, Cell Signaling Technolgy, 1:1000), PARP (9542, Cell Signalling Technology, 1:1000), ATGL (2138, Cell Signaling Technology, 1:1000) CPT1A (D3B3, 12252, Cell Signaling Technology, 1:1000), ACSL1 (9189, Cell Signaling Technology, 1:1000) and HIF2α (D9E3, 7096, Cell Signaling Technology, 1:1000) and beta-actin (AC-15, ab6276, Abcam, 1:6000) was used as loading control. Afterwards, membranes were incubated with appropriate horseradish peroxidase-linked secondary antibodies (111-036-045 and 115-036-062, Jackson ImmunoResearch, 1:12500) followed by detection of chemiluminescence with ECL Prime Western Blotting Detection Reagent (GEHealthcare) on a ChemiDoc Imaging System using Image Lab software (Bio-Rad Laboratories). ImageJ software was used for densitometric analysis of protein bands.

### Flow cytometric assessment of lipid peroxidation

200000, 66000 and 22000 cells (high, med and low) were seeded in 6-cm dishes which corresponds to 3000, 1000 and 333 cells in 96-wells. Ferroptosis in GPX4 knockout clones was induced by direct withdrawal of Lip1 from the culture medium and lipid peroxidation was measured the next day. Cells were stained with 2 µM BODIPY-C11 581/591 (ThermoFisher) for 30 min at 37°C. Cells were washed with PBS, harvested by trypsinization and resuspended in PBS containing 1% BSA for flow cytometry analysis. 1 µM Sytox blue staining (ThermoFisher) was used to discriminate live/dead cells. At least 10000 cells per sample were immediately recorded on a FACS Aria IIIu (BD Biosciences) using laser and filters according to manufacturer’s instructions. Data was analyzed using FlowJo Software (FlowJo, LLC).

### LipidTox staining of triacylglycerides

HMLE and HMLE-Twist1 cells were seeded at 3000, 1000 and 333 cells per well (high, med and low respectively) in 96-well plates with an optically clear bottom suitable for immunofluorescence (Perkin Elmer). After 48h, cells were fixed with 4% paraformaldehyde (PFA, Sigma) containing 1 mg/ml Hoechst 33342 (Thermo Fisher Scientifc) dye for nuclear staining and afterwards stained with LipidTOX™ Green Neutral Lipid Stain (Thermo Fisher Scientifc) according to manusfacturer’s instructions. Pictures were taken on a Zeiss Apoptome.2 (20x magnification) using ZEN imaging software.

### RNA preparation and RT-qPCR Analysis

For density experiments, 200000, 66000 and 22000 cells (high, med and low) were seeded in 6-cm dishes which corresponds to 3000, 1000 and 333 cells in 96-wells. Two days after plating, mRNA was isolated using the RNeasy Mini Kit (Qiagen) according to manufacturer’s instructions. 1 µg total RNA was reverse transcribed using Oligo(dT) primers for amplification (OneScript cDNA Kit, abm). For qPCR, specific primers were used in a power SYBR Green-PCR Master Mix reaction (Applied Biosystems) which was run on a Quantstudio 12K Flex qPCR System (Applied Biosystems). *RPL32* was used for normalization. The following primer sequences were used: *RPL32*: forward 5’-CAGGGTTCGTAGAAGATTCAAGGG-3’ and reverse 5’- CTTGGAGGAAACATTGTGAGCGATC-3’, ACSL1: forward 5’- CCATGAGCTGTTCCGGTATTT-3’ and reverse 5’- CCGAAGCCCATAAGCGTGTT-3’, ACSL3: forward 5’- GCCGAGTGGATGATAGCTGC-3’ and reverse 5’- ATGGCTGGACCTCCTAGAGTG-3’, ACSL4: forward 5’- ACTGGCCGACCTAAGGGAG-3’ and reverse 5’- GCCAAAGGCAAGTAGCCAATA-3’, SCD: forward 5’- TCTAGCTCCTATACCACCACCA-3’ and reverse 5’- TCGTCTCCAACTTATCTCCTCC- 3’, *CPT1A*: forward 5’-ATCAATCGGACTCTGGAAACGG-3’ and reverse 5’-TCAGGGAGTAGCGCATGGT-3’, *ACAA2*: forward 5’-CTGCTCCGAGGTGTGTTTGTA-3’ and reverse 5’-GGCAGCAAATTCAGACAAGTCA-3’

### CRISPR/Cas9-mediated knockout of individual genes

Benchling software as well as the MIT CRISPR design tool (http://crispr.mit.edu/) were used to design sgRNA guides for targeting critical exons of GPX4, ACSL4 and PTEN. For GPX4 and ACSL4 knockout, string assembly gRNA cloning (STAgR) was employed to clone sgRNAs into the STAgR_Neo plasmid as recently described ^50^. For Gibson Assembly reaction, a Gibson Assembly Master Mix (NEB) was used according to manufacturer’s instructions and diluted 1:4 prior to transformation into XL10-Gold ultracompetent cells (Agilent Technologies). For GPX4 and ACSL4 gene knockout, the CRISPR/Cas9 system was transiently expressed. Briefly, 150000 to 200000 cells were seeded in 6-well dishes and cultured for 24h. Cells were then co-transfected using TransIT-X2 transfection reagent (Mirus Bio LLC) at a ratio 1:3 (approximately 1 µg each) of a Cas9-GFP-expressing plasmid (pSpCas9(BB)-2A-GFP) and respective STAgR-Neo plasmid containing sgRNAs according to manufacturer’s instructions. 72h after transfection, cells were sorted for GFP expression on a FACSAriaIIIu (BD Biosciences), Single cells were seeded into 96-wells and expanded. To assess CRISPR/Cas9-induced deletions or insertions, PCRs of the expected corresponding genomic locus were performed and validated by sequencing and immunoblotting of the respective proteins. The following sgRNA sequences were used: GPX4: 5’-TTTCCGCCAAGGACATCGAC-3’, 5’-CGTGTGCATCGTCACCAACG-3’ and 5’-ACTCAGCGTATCGGGCGTGC-3’, ACSL4: 5’-ATTGTTATTAACAAGTGGAC-3’, 5’-CTAGCTGTAATAGACATCCC-3’ and 5’- TGCAATCATCCATTCGGCCC-3’. For PTEN deletion, sgRNA was cloned into a pLX-sgRNA vector (Addgene plasmid#50662) ^51^. HMLE-Twist1-ER 24^hi^ cells were stably transduced with lentiviruses expressing a doxycycline-inducible Cas9 protein (pCW-Cas9, Addgene #50661) ^51^ and selected with 1 µg/ml puromycin (Sigma). Then, cells were stably transduced with lentiviruses containing the sgRNAs against PTEN. In these cells, Cas9 expression was activated with 0.5 µg/ml doxycycline treatment (Sigma) to allow CRISPR/Cas9-mediated modification in the PTEN locus and single cells were seeded in 96-well plates. Clones were screened by performing PCR and sequencing of the expected corresponding genomic locus. Successful PTEN deletion was validated by assessing activation status of downstream AKT signaling by immunoblotting. The following sgRNA sequence was used to target PTEN: 5’-TGTGCATATTTATTACATCG-3’.

### siRNA mediated gene knockdown

2x 10^5^ cells were seeded in 6-wells and the next day transfected with 25 nM SMARTpool siGENOME human siRNAs targeting genes of interest (ACSL1, ACSL3, ACSL4, Dharmacon) using TransIT-X2 transfection reagent (Mirus Bio LLC) according to manufacturer’s instructions. Transfections with siGENOME Non-Targeting Pool #2 siRNAs (si-nt, Dharmacon) and untransfected cells were used as controls. 24h after transfection, cells were used for further experiments and 72h after transfection, knockdown efficiency was assessed using RT-qPCR.

### Culture in 3D collagen gels, carmine staining of gels and 3D immunofluorescence

3D collagen gels (final concentration 1.3 mg/ml collagen I) were prepared and cultured as previously described ^26^. For HMLE SCC H5 *GPX4* knockout, 675, 225 and 75 cells (high, med and low) per 24-well gel were seeded in 500 nM Lip1 containing medium or directly withdrawn of Lip1. After 3 days, all gels were washed with PBS and for the +/− Lip1 condition, Lip1 was removed by changing the culture medium. For primary mammary epithelial cell culture, 5000 cells were seeded per 24-well gel. After 5 days of initial survival culture in mammary epithelial cell growth medium (MECGM, Promocell) containing 3 µM Y-27632 (Biomol), 10 µM Forskolin (Adipogen) and 0.5% FCS (PAN Biotech), MECGM medium was supplemented with a final concentration of 0.1% DMSO, 100 nM RSL3, 500 nM Lip1, or the combination of 100 nM RSL3 and 500 nM Lip1. Medium was replaced every 2 days and gels were fixed with 4% paraformaldehyde (PFA, Sigma) after 7-11 days of culture. Carmine staining and imaging was performed as previously described ^26^. Pictures were analyzed by ImageJ software. For this purpose, images were first converted to a binary image and colonies extracted from the background by setting a manual threshold. Particles with an area between 400 and 90000 µm^2^ (20-300 µM diameter) and a circularity 0.5-1 were counted. 3D immunofluorescence was performed as previously described ^26^. Primary antibodies used were: Vimentin (V9, MAB3578, Abnova, 1:100) and Ki67 (ab15580, Abcam, 1:300). Following secondary antibodies used were: Donkey anti-Rabbit IgG Alexa Fluor 546 (A10040, Thermo Fisher, 1:250), Donkey anti-Mouse IgG Alexa Fluor 488 (A21202, Thermo Fisher, 1:250). Pictures were taken on an Olympus Confocal (20x magnification) using FV10-ASW software.

### Proteomic analysis of cells seeded in cell density

516000, 172000 and 57000 cells (high, med and low) were seeded on 10-cm dishes, corresponding to 3000, 1000 and 333 cells in 96-wells. Proteins of cells were isolated 48h after seeding. Protein isolation and concentration measurement was performed as described in immunoblotting section.

### Proteomic MS sample preparation

10 µg of sample were enzymatically digested using a modified filter-aided sample preparation (FASP) protocol ^52, 53^. Peptides were stored at -20°C until MS measurement.

### Proteomic MS measurement

Mass spectrometry (MS) measurements were performed in data dependent (DDA) mode. MS data were acquired on a Q Exactive (QE) high field (HF) mass spectrometer (Thermo Fisher Scientific Inc.). Approximately 0.5 μg per sample were automatically loaded to the online coupled RSLC (Ultimate 3000, Thermo Fisher Scientific Inc.) HPLC system. A nano trap column was used (300 μm inner diameter (ID) × 5 mm, packed with Acclaim PepMap100 C18, 5 μm, 100 Å; LC Packings, Sunnyvale, CA) before separation by reversed phase chromatography (Acquity UPLC M-Class HSS T3 Column 75µm ID x 250mm, 1.8µm; Waters, Eschborn, Germany) at 40°C. Peptides were eluted from column at 250 nl/min using increasing acetonitrile (ACN) concentration (in 0.1% formic acid) from 3% to 41 % over a 105 minutes gradient. The high-resolution (60 000 full width at half-maximum) MS spectrum was acquired with a mass range from 300 to 1500 m/z with automatic gain control target set to 3 x 10^6^ and a maximum of 50 ms injection time. From the MS prescan, the 10 most abundant peptide ions were selected for fragmentation (MSMS) if at least doubly charged, with a dynamic exclusion of 30 seconds. MSMS spectra were recorded at 15 000 resolution with automatic gain control target set to 1 x 10^5^ and a maximum of 100 ms injection time. Normalized collision energy was set to 28 and all spectra were recorded in profile type.

### Progenesis QI for label-free quantification for proteomics

Spectra were analyzed using Progenesis QI software for proteomics (Version 3.0, Nonlinear Dynamics, Waters, Newcastle upon Tyne, U.K.) for label-free quantification as previously described ^53^ with the following changes: spectra were searched against the Swissprot human database (Release 2017.02, 553473 sequences). Search parameters used were 10 ppm peptide mass tolerance and 20 mmu fragment mass tolerance. Carbamidomethylation of cysteine was set as fixed modification and oxidation of methionine and deamidation of asparagine and glutamine was allowed as variable modifications, allowing only one missed cleavage site. Mascot integrated decoy database search was set to a false discovery rate (FDR) of 1 %.

### Analysis of proteomic study

For each cell line, ratios of the normalized, averaged protein abundance (n=4) of the high cell density condition (set to 1) and the respective averaged abundance of the low cell density condition were calculated. Ratios >1 indicated enriched proteins at low cell densities and ratios <1 indicated enriched proteins at high cell density. To assess significantly, regulated proteins, an unpaired, two-tailed T-test with Welch’s correction was performed on log2 expression data of high cell density compared to low cell density within each cell lines. Proteins with a p-value below 0.05 were considered to be significantly regulated. The overlap of significantly, regulated proteins in the same direction shared by both cell lines (irrespective of fold changes) were submitted to GO term enrichment analysis using DAVID ^34, 35^. Potential hits were further analyzed by functional assays.

### Lipidomic analysis of cells seeded in cell density

516000, 172000 and 57000 cells (high, med and low) were seeded on 10-cm dishes, corresponding to 3000, 1000 and 333 cells in 96-wells and cultured for 48h. Cells were harvested by trypsinization, washed with PBS and pelleted by centrifugation. After aspiration of supernatant, cell pellets were snap frozen in liquid nitrogen and stored at -80°C until lipidomic analysis.

### Lipid extraction for lipidomics

700 μl of sample were mixed with 800 μl 1 N HCl:CH3OH 1:8 (v/v), 900 μl CHCl3 and 200 μg/ml of the antioxidant 2,6-di-tert-butyl-4-methylphenol (BHT; Sigma Aldrich). 3 μl of SPLASH® LIPIDOMIX® Mass Spec Standard (#330707, Avanti Polar Lipids) was spiked into the extract mix. The organic fraction was evaporated using a Savant Speedvac spd111v (Thermo Fisher Scientific) at room temperature and the remaining lipid pellet was stored at -20°C under argon.

### Lipidomic mass spectrometry

Just before mass spectrometry analysis, lipid pellets were reconstituted in 100% ethanol. Lipid species were analyzed by liquid chromatography electrospray ionization tandem mass spectrometry (LC-ESI/MS/MS) on a Nexera X2 UHPLC system (Shimadzu) coupled with hybrid triple quadrupole/linear ion trap mass spectrometer (6500+ QTRAP system; AB SCIEX). Chromatographic separation was performed on a XBridge amide column (150 mm × 4.6 mm, 3.5 μm; Waters) maintained at 35°C using mobile phase A [1 mM ammonium acetate in water-acetonitrile 5:95 (v/v)] and mobile phase B [1 mM ammonium acetate in water-acetonitrile 50:50 (v/v)] in the following gradient: (0-6 min: 0% B to 6% B; 6-10 min: 6% B to 25% B; 10-11 min: 25% B to 98% B; 11-13 min: 98% B to 100% B; 13-19 min: 100% B; 19-24 min: 0% B) at a flow rate of 0.7 mL/min which was increased to 1.5 mL/min from 13 minutes onwards. Sphingomyelins (SM), d18:1 and d18:0 ceramides (Cer), hexosylceramides (HexCer), lactosylceramides (LacCer) were measured in positive ion mode with a precursor scan of 184.1, 264.4 and 266.4, 264.4 and 264.4 respectively. Triacylglycerides (TAG) and diacylglycerides (DAG) were measured in positive ion mode with a neutral loss scan for one of the fatty acyl moieties. Phosphatidylcholine (PC), lysophosphatidylcholine (LPC), phosphatidylethanolamine (PE), lysophosphatidyl-ethanolamine (LPE), phosphatidylglycerol (PG), phosphatidylinositol (PI) and phosphatidylserine (PS) were measured in negative ion mode with a neutral loss scan for the fatty acyl moieties. Lipid quantification was performed by scheduled multiple reactions monitoring (MRM), the transitions being based on the neutral losses or the typical product ions as described above. The instrument parameters were as follows: Curtain Gas = 35 psi; Collision Gas = 8 a.u. (medium); IonSpray Voltage = 5500 V and −4,500 V; Temperature = 550°C; Ion Source Gas 1 = 50 psi; Ion Source Gas 2 = 60 psi; Declustering Potential = 60 V and −80 V; Entrance Potential = 10 V and −10 V; Collision Cell Exit Potential = 15 V and −15 V.

### Data analysis Lipidomics

Peak integration was performed with the MultiQuantTM software version 3.0.3. Lipid species were normalized to the internal standards and corrected for natural isotope abundance. Isotopic mass distributions were calculated with Python Molmass 2019.1.1. Unpaired T-test p-values and FDR corrected p-values (using the Benjamini/Hochberg procedure) were calculated with Python StatsModels version 0.10.1.

### 13C_6_-glucose metabolomics of cell seeded in cell density

516000, 172000 and 57000 cells (high, med and low) were seeded on 10-cm dishes, corresponding to 3000, 1000 and 333 cells in 96-wells. After attachment, cells were washed once with 0.9% saline (pH 7.4) and cultured for 48h in mammary epithelial growth medium (Promocell, ordered in custom formulation without glucose) supplemented with 8 mM ^13^C_6_-glucose (Sigma). After 48h, cells were washed with 0.9% saline (pH 7.4) and harvested by scraping cells in 0.9% saline prior to pelleting cells by centrifugation. After aspiration of supernatant, cell pellets were snap frozen in liquid nitrogen and stored at -80°C until metabolomic analysis.

### Metabolites extraction and derivatization method for metabolomics

Metabolite extraction were previously described and must be performed in a mixture of dry and wet ice ^54, 55^. Briefly, a volume of 800 µL of methanol/water (60/40) (v/v) containing 90 ng/ml of glutaric acid as internal standard was used for the extraction of metabolites. Each sample was transferred in an Eppendorf tube and 500 µL of chloroform containing C17 internal standard at a concentration of 10 µg/ml was subsequently added. Samples were vortexed at 4°C during 10 min and centrifuged at the maximum speed for 10 min at 4°C. Metabolites were divided in two phases separated by a protein layer: polar metabolites in the methanol/water (upper) phase and the lipid fraction in the chloroform (lower) phase. Following metabolites separation, every phase was dried at 4°C overnight using a vacuum concentration.

The samples were derivatized and measured as described before ^56^. Briefly, fatty acids were esterified with 500 µL of 2 % sulfuric acid in methanol and incubated overnight at 50 °C. Subsequently, fatty acids were extracted with 600 µL of MS-grade hexane and 100 µL of saturated NaCl solution. Hexane fraction was dried in a vacuum concentration at room temperature for 30 min and was resuspended in 50 µL of hexane before to be analyzed by gas chromatography coupled to mass spectrometry (GC-MS).

### Metabolomic gas chromatography–mass spectrometric analysis

The fatty acids were analyzed by gas chromatography (7890A GC system) coupled to mass spectrometry (5975C Inert MS system) from Agilent Technologies. The metabolite separation was performed with a DB35MS column (30 m, 0.25mm, 0.25 µm) from Agilent Technologies using a carrier gas flow of helium fixed at 1.3 ml/min. The inlet temperature was set at 270 °C and 1 µL of sample were injected with a split ratio 1 to 3. To ensure the separation of fatty acids, the initial gradient temperature was set at 140 °C for 2 min and increased at the ramping rate of 1°C/min to 185°C, following by a ramping rate of 20°C/min to rich 300°C. The temperatures of the quadrupole and the source were set at 150°C and 230°C, respectively. The electron impact ionization was set at 70 eV and a full scan mode ranging from 100 to 600 a.m.u (mass) was applied for the detection of fatty acids.

### Metabolomic data analysis – Matlab

The results were analyzed using an in-house Matlab script to extract mass distribution vectors and to integrate raw ion chromatograms. The natural isotopes distribution were corrected using the method developed by Fernandez et al, 1996 ^57^. For each sample, the area of fatty acid metabolites was subsequently normalized to the protein content and to the C17 internal standard area. With the setup used, cis-C18:1 isomers like oleate and cis-8-octadecenoate cannot be separated ^58^. However, since cis-8-octadecenoate is an elongation product of sapienate and sapienate is very low ^58^, we assumed that the cis8/9-C18:1 peak only represents oleate. A Matlab M-file was also used to perform an isotopomer spectral analysis (ISA) ^59^ to determine the fraction of newly synthesized palmitate, stearate and oleate.

### Data presentation and statistical analyses

Data are presented as mean ± s.e.m. of n = x experiments, with x indicating the number of experiments independently performed, unless stated otherwise. Statistical analysis was performed using GraphPad Prism 8.1.2 software or Excel 2016. In general, unless stated otherwise, an unpaired, two-tailed T-test was performed with Welch’s correction and a p-value below 0.05 was considered significant.

## Acknowledgements

We thank Christopher Breunig and Stefan Stricker (Institute of Stem Cell Research, Helmholtz Zentrum Munich) for sharing reagents and guidance for STAgR cloning. We thank members of the Scheel Group, especially Lisa Meixner and Laura Eichelberger, for sharing experimental expertise. Further, we thank Thomas Schwarz-Romond, Alecia-Jane Twigger, Massimo Saini and Laura Eichelberger for critical reading of the manuscript, Magdalena Götz, Andreas Jung and all members of the Scheel group for productive discussion of the data. E.P. was supported by a Boehringer Ingelheim Fonds PhD fellowship. S.-M.F. acknowledges funding from the European Research Council under the ERC Consolidator Grant Agreement n. 771486–MetaRegulation and from FWO projects and KU Leuven Methusalem Co-funding. This work has been supported by the Interreg VA EMR program (EURLIPIDS, EMR23) (J.V.S.). This work has been further supported by a grant of the German Cancer Aid Foundation (Max Eder Program, Deutsche Krebshilfe 110225) to C.H.S., and by grants by the German Federal Ministry of Education and Research (BMBF) through the Joint Project Modelling ALS Disease In Vitro (MAIV, 01EK1611B) and the VIP + program NEUROPROTEKT (03VP04260) to M.C.

## Author’s contributions

E.P. and C.H.S. conceived the study and designed experiments. E.P., F.H., M.B.-H. and H.M.G. performed *in vitro* experiments and analyzed data. C.v.T., A.-C.K. and S.M.H. performed proteomics and analyzed the data. J.D., A.T. and J.V.S. performed lipidomics and analyzed the data. M.P. and S.-M.F. performed ^13^C_6_ glucose metabolomics and analyzed the data. S.D. and J.P.F.A. provided reagents and participated in discussion, evaluation and interpretation of the data. E.P. assembled the figures. E.P., M.C. and C.H.S. interpreted the data and wrote the manuscript. All authors read and approved the final manuscript.

---

**Supplementary Fig. 1.**
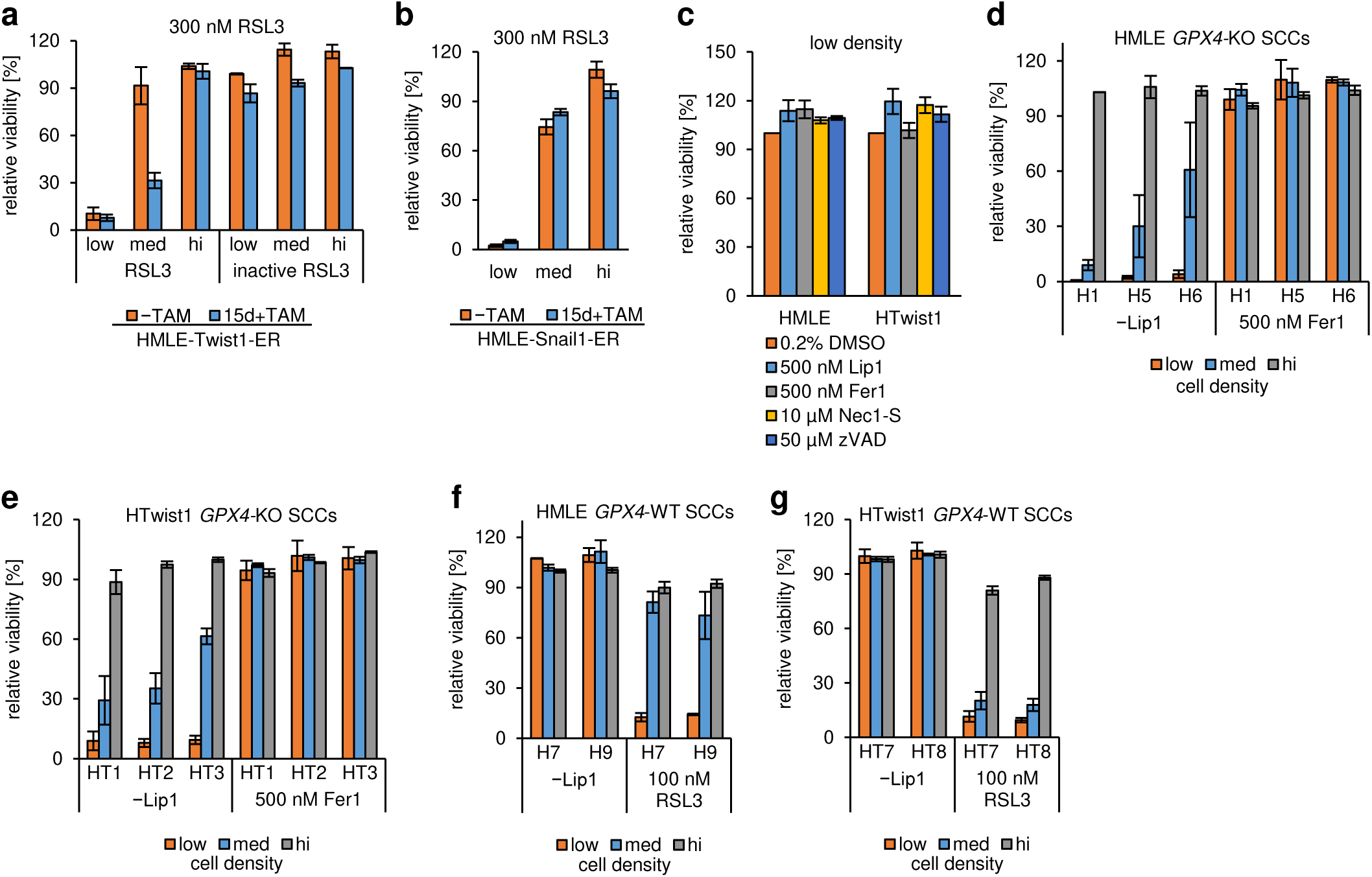
Cell density sensitizes mammary epithelial cells to ferroptosis irrespectively of cellular phenotype. **a** Viability assay: treatment of epithelial (−TAM) or 15 days 4-hydroxytamoxifen-induced mesenchymal (15d+TAM) HMLE-Twist1-ER 24^high^ cells with 0.3% DMSO control, 300 nM RSL3 or 300 nM of an inactive isomer of RSL3 in density, n=3. **b** Viability assay: treatment of epithelial (−TAM) or 15 days 4-hydroxytamoxifen-induced mesenchymal (15d+TAM) HMLE-Snail1-ER 24^high^ cells with 0.3% DMSO control or 300 nM RSL3, n=1. **c** Rescue-viability assay: control treatment HMLE and HMLE-Twist1 cells with DMSO, 500 nM ferrostatin1 (Fer1), 500 nM liproxstatin1 (Lip1), 10 µM necrostatin1-S (Nec1-S) or 50 µM zVAD-fmk (zVAD) at low density, mean of at least three biological replicates is shown (n=3-5), related to Fig. 1e. **d** Viability assay: CRISPR/Cas9-derived single cell clones (SCCs) of HMLE upon 1 µM Lip1, Lip1 withdrawal or 500 nM Fer1 at density, n=2, related to Fig.1g, h. **e** Viability assay: CRISPR/Cas9-derived HMLE-Twist1 SCCs as described in d, n=3-4, related to Fig.1g, h. **f** Viability assay of CRISPR/Cas9-derived control HMLE SCCs with wildtype (WT) GPX4 expression upon 1 µM Lip1 (control), 100 nM RSL3 or deprivation of Lip1 at density, n=2, related to Fig. 1g, h. **g** Viability assay of CRISPR/Cas9-derived HMLE-Twist1 SCCs as described in f, n=2-4, related to Fig. 1g, h. Data are presented as mean of indicated biological replicates ± s.e.m. (n=x). Data was normalized to respective DMSO control (**a-c**) or to respective Lip1 control (**d-g**) within each seeding density and cell line. Statistics: two-tailed, unpaired T-test with Welch’s correction (p-value: *<0.05, **<0.01, ***0.001, ****<0.0001, n.s. = not significant).

**Supplementary Fig. 2.**
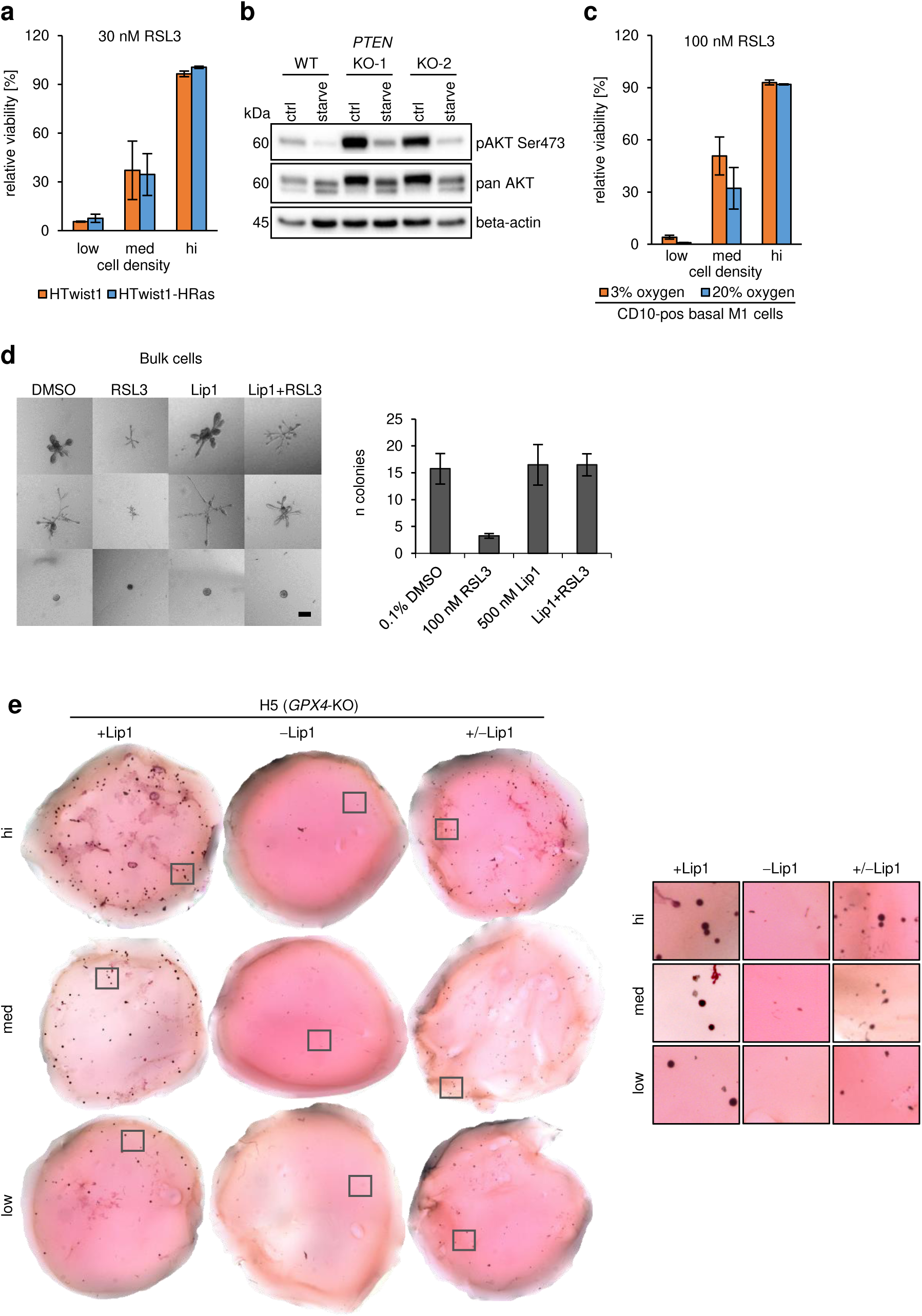
Cell density-dependent cell death is maintained during cellular transformation and prevents growth in 3D-collagen gels. **a** Viability assay: treatment of HMLE-Twist1 and HMLE-Twist1-HRas with 0.3% DMSO or 30 nM RSL3 cells in density, n=2. **b** Immunoblot: phosphorylated AKT at serine residue 473 (pAKT Ser473), total AKT 1 and 2 isoforms (pan AKT) and beta-actin protein expression in *PTEN*-wildtype (WT) and two *PTEN*-knockout clones (KO-1 and KO-2) of HMLE-Twist1-ER cells (without Twist1-activation). Cells were grown in supplemented mammary epithelial growth medium (ctrl) or in basal DMEM/F-12 medium (starve). beta-actin serves as loading control. kDa = kilo Dalton. **c** Viability assay: treatment of sorted, CD10-positive primary mammary epithelial cells of the basal lineage (B cells) of Donor M1 with 0.1% DMSO or 100 nM RSL3 at ambient oxygen level (20%) or oxygen levels present in tissues (normoxia, 3%) in density, n=3. **d** 3D-collagen gels: Bulk primary mammary epithelial cells were treated with 0.1% DMSO, 100 nM RSL3, 500 nM Lip1 or a combination of RSL3 and Lip1 for 7-10d prior quantification of arising colonies. Bright-field images of representative colonies and the mean of 3-4 technical replicates ± s.d. of one experiment performed is shown. Scale bar: 200 µm. **e** 3D-collagen gels: representative carmine stainings (left) with magnifications (right) showing colonies in black of HMLE single cell clone H5 with *GPX4*-knockout (KO) are shown. H5 was seeded at indicated densities in 3D collagen gels in medium containing 500 nM Lip1 (+Lip1), withdrawn of Lip1 either directly (-Lip1) or after 3d in initial +Lip1 culture (+/-Lip1) for 7-10 days, related to Fig. 2g. Data (a, c) are presented as mean of indicated biological replicates ± s.e.m. (n=x) and were normalized to respective DMSO control within each density and cell line. Statistics: two-tailed, unpaired T-test with Welch’s correction (p-value: *<0.05, **<0.01, ***0.001, ****<0.0001, n.s. = not significant).

**Supplementary Fig. 3.**
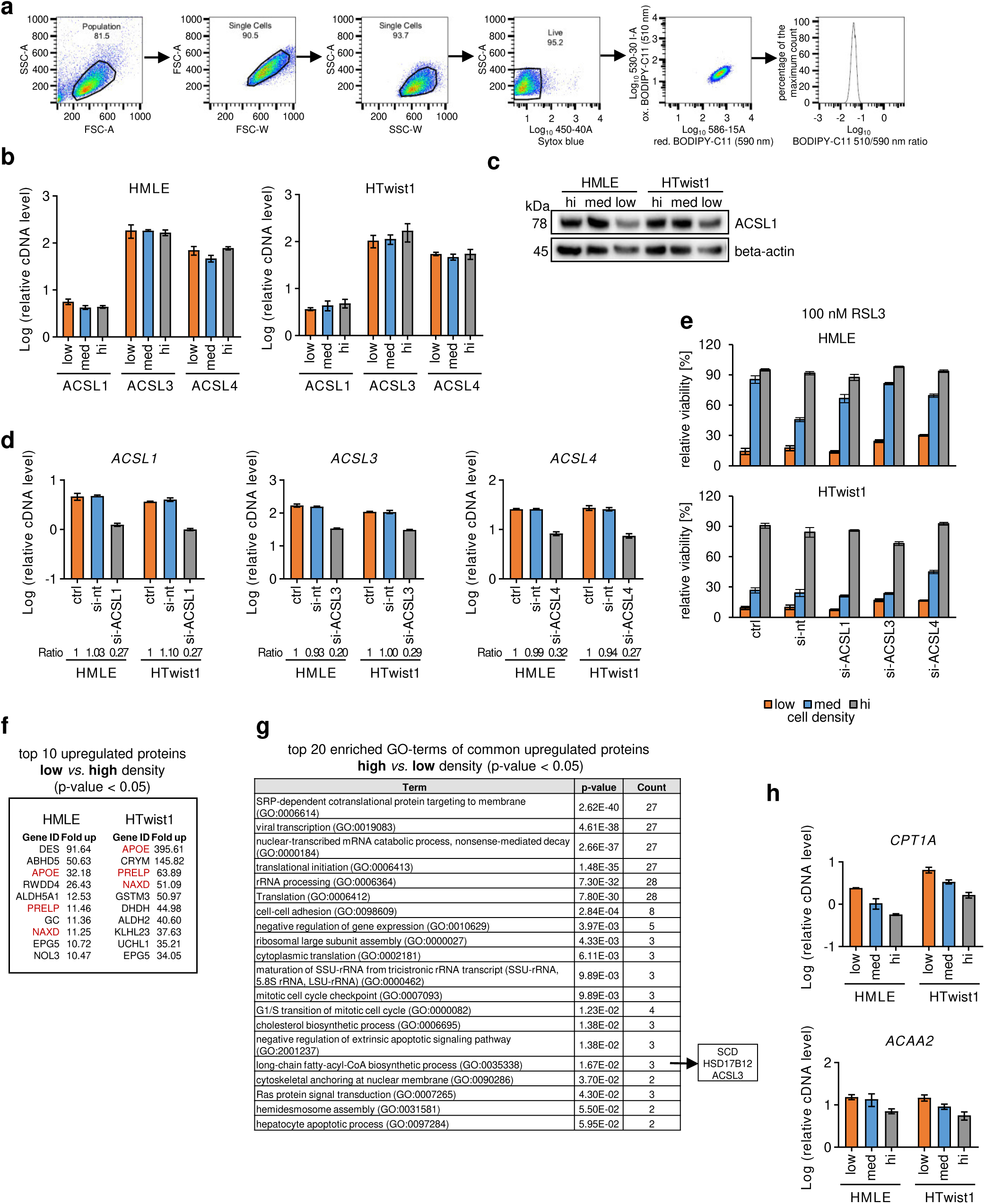
Cell density-dependent ferroptosis does not show all classical hallmarks of ferroptosis and is independent of ACSL enzymes. **a** Flow cytometry: Example gating strategy for BODIPY-C11 staining of *GPX4*-knockout SCCs as described in Fig. 3b. Ox: oxidized BODIPY-C11 (510 nm), Red: reduced BODIPY-C11 (590 nm). **b** RT-qPCR: *ACSL1*, *ACSL3 and ACSL4* mRNA expression of HMLE and HMLE-Twist1 cells seeded in density. *RPL32* was used as an internal control. Mean of three technical replicates ± s.e.m. of one representative experiment from two independently performed experiments is shown. Y-axis: log10. **c** Immunoblot: ACSL1 and beta-actin (loading control) protein expression in HMLE and HMLE-Twist1 cells seeded in density. kDa= kilo Dalton. **d** RT-qPCR: *ACSL1*, *ACSL3 and ACSL4* mRNA expression of HMLE and HMLE-Twist1 cells at intermediate seeding density 72h after transfection with non-targeting (si-nt) or siRNA pools targeting *ACSL1*, *ACSL3* or *ACSL4*. Untransfected control (ctrl) is shown. Ratios represent knockdown efficiency normalized to ctrl. *RPL32* was used as an internal control. Mean of three technical replicates ± s.e.m is shown. **e** Viability assay: treatment of HMLE and HMLE-Twist1 cells transfected with siRNAs as described in Supplementary Fig. 3d with 100 nM RSL3 for 24h. Data was normalized to respective DMSO control within each cell line and density. Mean of three technical replicates ± s.e.m is shown of one representative experiment performed two times. **f** Proteomic analysis: top 10, significantly enriched proteins at low cell density compared to high cell density for each cell line of the commonly, upregulated data set at low cell density (143 proteins), related to Fig. 3i, n=4. g GO-term analyses: top 20, significantly enriched terms (biological processes) of commonly, upregulated proteins (89) in high cell density compared to low cell density in both HMLE and HMLE-Twist1 cells. Count indicates the number of genes (proteins) of the data set found within the term, related to Fig. 3h, n=4. h RT-qPCR: *CPT1A* and *ACAA2* mRNA expression of HMLE and HMLE-Twist1 cells seeded in density. *RPL32* was used as an internal control. Mean of three technical replicates ± s.e.m. of one representative experiment from two independently performed experiments is shown. Y-axis: log10.

**Supplementary Fig. 4.**
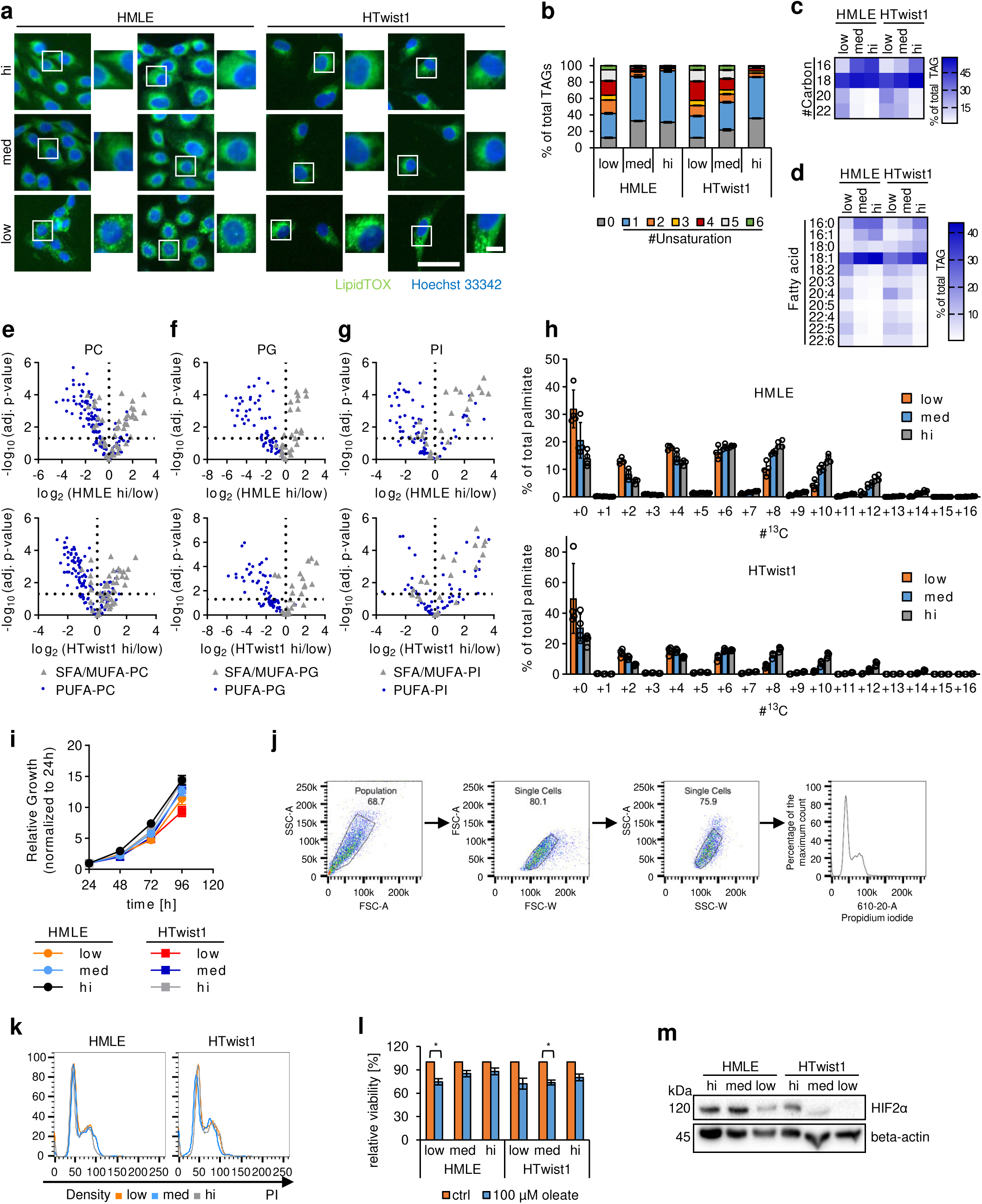
Epithelial and Twist1-induced mesenchymal HMLE cells enrich polyunsaturated triacylglycerides at low cell density. **a Immunofluorescence:** representative images of neutral triacylglycerides/lipid droplet staining using LipidTOX (green) in HMLE and HMLE-Twist1 cells seeded at density. Cell nuclei were visualized using Hoechst 33342 staining (blue). Scale bar: 50 µm. Scale bar of indicated blow ups: 10 µm. **b** TAG species summed by the same number of unsaturation shown as percentage of acyl chains containing the respective unsaturation within each cell line and density, n=4, related to Fig. 4d. **c** TAG species summed by the same number of chain carbon length shown as percentage of acyl chains containing the respective chain length in a heatmap within each cell line and density, n=4, related to Fig. 4d. **d** Relative fatty acid composition of TAGs shown as percentage within each cell line and density, n=4, related to Fig. 4d. **e** Volcano plot: log2 transformed fold changes of relative PC species abundance between high (hi) and low cell density for each cell line plotted against -log10 FDR adjusted p-values calculated using the Benjamini–Hochberg procedure. PC species were divided into SFA/MUFA-PCs (grey triangle) or PUFA-PCs (blue dot). n=4. **f** Volcano plot: fold changes of relative PG species abundance grouped as SFA/MUFA-PGs (grey triangle) or PUFA-PGs (blue dot) between high (hi) and low cell density for each cell line, plotted as described in Supplementary Fig. 4e. n=4. **g** Volcano plot: fold changes of relative PI species abundance grouped as SFA/MUFA-PIs (grey triangle) or PUFA-PIs (blue dot) between high (hi) and low cell density for each cell line, plotted as described in Supplementary Fig. 4e. n=4. **h** Mass isotopomer distribution of palmitate in HMLE and HMLE-Twist1 seeded at density and incubated for 48h with media containing 8 mM ^13^C_6_ glucose. n=4. **i** Proliferation curve: measurement of metabolically-active HMLE and HMLE-Twist1 cells plated at indicated densities in 96-wells every 24h for a period of 96h. Data shown represent mean ± s.d. of n=10 wells per time point normalized to the 24h measurement. **j** Flow cytometry: Example gating strategy for cell cycle analysis using propidium iodide staining of HMLE and HMLE-Twist1 cells seeded at different densities. Related to Supplementary Fig. 4k. **k** Cell-cycle analysis by flow cytometry: staining of HMLE and HMLE-Twist1 cells seeded at indicated densities in 10-cm dishes for 48h prior to fixation and staining with propidium iodide (PI). X-axis: linear scale of PI fluorescence. Y-axis: percentage of maximum count. Data show one representative experiment performed independently two times. **l** Viability assay: treatment of HMLE and HMLE-Twist1 seeded at indicated densities with 0.1% DMSO only (ctrl, orange) or pre-treated for 24h with 100 µM oleate (blue). Oleate was present during DMSO treatment, n=3, relates to Fig. 4k. **m** Immunoblot: HIF2α and beta-actin (loading control) protein expression in HMLE and HMLE-Twist1 cells seeded in density, kDa = kilo Dalton. Data are presented as mean of indicated biological replicates ± s.e.m. (**b, l**) or ± s.d. (**h, i**).

## References

1. Dixon, S. J. et al. Ferroptosis: an iron-dependent form of nonapoptotic cell death. Cell 149, 1060–72 (2012).

2. Stockwell, B. R. et al. Ferroptosis: A Regulated Cell Death Nexus Linking Metabolism, Redox Biology, and Disease. Cell 171, 273–285 (2017).

3. Conrad, M., Angeli, J. P. F., Vandenabeele, P. & Stockwell, B. R. Regulated necrosis: disease relevance and therapeutic opportunities. Nat. Rev. Drug Discov. 15, 348–66 (2016).

4. Friedmann Angeli, J. P., Krysko, D. V. & Conrad, M. Ferroptosis at the crossroads of cancer-acquired drug resistance and immune evasion. Nat. Rev. Cancer 19, 405–414 (2019).

5. Tsoi, J. et al. Multi-stage Differentiation Defines Melanoma Subtypes with Differential Vulnerability to Drug-Induced Iron-Dependent Oxidative Stress. Cancer Cell 33, 890–904.e5 (2018).

6. Viswanathan, V. S. et al. Dependency of a therapy-resistant state of cancer cells on a lipid peroxidase pathway. Nature 547, 453–457 (2017).

7. Hangauer, M. J. et al. Drug-tolerant persister cancer cells are vulnerable to GPX4 inhibition. Nature 551, 247–250 (2017).

8. Yang, W. S. et al. Regulation of ferroptotic cancer cell death by GPX4. Cell 156, 317–331 (2014).

9. Seiler, A. et al. Glutathione peroxidase 4 senses and translates oxidative stress into 12/15-lipoxygenase dependent- and AIF-mediated cell death. Cell Metab. 8, 237–48 (2008).

10. Friedmann Angeli, J. P. et al. Inactivation of the ferroptosis regulator Gpx4 triggers acute renal failure in mice. Nat. Cell Biol. 16, 1180–91 (2014).

11. Ishii, T., Sugita, Y. & Bannai, S. Regulation of glutathione levels in mouse spleen lymphocytes by transport of cysteine. J. Cell. Physiol. 133, 330–336 (1987).

12. Hayano, M., Yang, W. S., Corn, C. K., Pagano, N. C. & Stockwell, B. R. Loss of cysteinyl-tRNA synthetase (CARS) induces the transsulfuration pathway and inhibits ferroptosis induced by cystine deprivation. Cell Death Differ. 23, 270–278 (2016).

13. Doll, S. et al. ACSL4 dictates ferroptosis sensitivity by shaping cellular lipid composition. Nat. Chem. Biol. 13, 91–98 (2017).

14. Kagan, V. E. et al. Oxidized arachidonic and adrenic PEs navigate cells to ferroptosis. Nat. Chem. Biol. 13, 81–90 (2017).

15. Dixon, S. J. et al. Human Haploid Cell Genetics Reveals Roles for Lipid Metabolism Genes in Nonapoptotic Cell Death. ACS Chem. Biol. 10, 1604–1609 (2015).

16. Nieto, M. A., Huang, R. Y. Y. J., Jackson, R. A. A. & Thiery, J. P. P. Emt: 2016. Cell 166, 21–45 (2016).

17. Ansieau, S., Collin, G. & Hill, L. EMT or EMT-Promoting Transcription Factors, Where to Focus the Light? Front. Oncol. 4, 353 (2014).

18. Mani, S. a et al. The epithelial-mesenchymal transition generates cells with properties of stem cells. Cell 133, 704–15 (2008).

19. Yang, J. et al. Twist, a Master Regulator of Morphogenesis, Plays an Essential Role in Tumor Metastasis Ben Gurion University of the Negev. Cell 117, 927–939 (2004).

20. Casas, E. et al. Snail2 is an essential mediator of twist1-induced epithelial mesenchymal transition and metastasis. Cancer Res. 71, 245–254 (2011).

21. Schmidt, J. M. et al. Stem-Cell-like Properties and Epithelial Plasticity Arise as Stable Traits after Transient Twist1 Activation. Cell Rep. 10, 131–139 (2015).

22. Slee, E. A. et al. Benzyloxycarbonyl-Val-Ala-Asp (OMe) fluoromethylketone (Z-VAD.FMK) inhibits apoptosis by blocking the processing of CPP32. Biochem. J. 315, 21–24 (1996).

23. Degterev, A. et al. Chemical inhibitor of nonapoptotic cell death with therapeutic potential for ischemic brain injury. Nat. Chem. Biol. 1, 112–119 (2005).

24. Boulares, a H. et al. Role of Poly (ADP-ribose) Polymerase (PARP) Cleavage in Apoptosis. J. Biol. Chem. 274, 22932–22940 (1999).

25. Wolf, B. B., Schuler, M., Echeverri, F. & Green, D. R. Caspase-3 is the primary activator of apoptotic DNA fragmentation via DNA fragmentation factor-45/inhibitor of caspase-activated DNase inactivation. J. Biol. Chem. 274, 30651–30656 (1999).

26. Linnemann, J. R. et al. Quantification of regenerative potential in primary human mammary epithelial cells. Development 142, 3239–3251 (2015).

27. Brown, C. W., Amante, J. J., Goel, H. L. & Mercurio, A. M. The α6β4 integrin promotes resistance to ferroptosis. J. Cell Biol. 216, 4287–4297 (2017).

28. Schneider, M. et al. Absence of Glutathione Peroxidase 4 Affects Tumor Angiogenesis through Increased 12/15-Lipoxygenase Activity. Neoplasia 12, 254– 263 (2010).

29. Yang, W. S. et al. Peroxidation of polyunsaturated fatty acids by lipoxygenases drives ferroptosis. Proc. Natl. Acad. Sci. 113, E4966–E4975 (2016).

30. Drummen, G. P. C., Van Liebergen, L. C. M., Op den Kamp, J. A. F. & Post, J. A. C11-BODIPY581/591, an oxidation-sensitive fluorescent lipid peroxidation probe: (Micro)spectroscopic characterization and validation of methodology. Free Radic. Biol. Med. 33, 473–490 (2002).

31. Cao, J. Y. & Dixon, S. J. Mechanisms of ferroptosis. Cell. Mol. Life Sci. 73, 2195– 2209 (2016).

32. Kim, J. H., Lewin, T. M. & Coleman, R. A. Expression and characterization of recombinant rat acyl-CoA synthetases 1, 4, and 5: Selective inhibition by triacsin C and thiazolidinediones. J. Biol. Chem. 276, 24667–24673 (2001).

33. Magtanong, L. et al. Exogenous Monounsaturated Fatty Acids Promote a Ferroptosis-Resistant Cell State. Cell Chem. Biol. 26, 420–432.e9 (2019).

34. Huang, D. W., Sherman, B. T. & Lempicki, R. A. Systematic and integrative analysis of large gene lists using DAVID bioinformatics resources. Nat. Protoc. 4, 44–57 (2009).

35. Huang, D. W., Sherman, B. T. & Lempicki, R. A. Bioinformatics enrichment tools: Paths toward the comprehensive functional analysis of large gene lists. Nucleic Acids Res. 37, 1–13 (2009).

36. Röhrig, F. & Schulze, A. The multifaceted roles of fatty acid synthesis in cancer. Nat. Rev. Cancer 16, 732–749 (2016).

37. Wang, H. et al. Unique regulation of adipose triglyceride lipase (ATGL) by perilipin 5, a lipid droplet-associated protein. J. Biol. Chem. 286, 15707–15715 (2011).

38. Smirnova, E. et al. ATGL has a key role in lipid droplet/adiposome degradation in mammalian cells. EMBO Rep. 7, 106–113 (2006).

39. Currie, E., Schulze, A., Zechner, R., Walther, T. C. & Farese, R. V. Cellular fatty acid metabolism and cancer. Cell Metab. 18, 153–161 (2013).

40. Jarc, E. et al. Lipid droplets induced by secreted phospholipase A2 and unsaturated fatty acids protect breast cancer cells from nutrient and lipotoxic stress. Biochim. Biophys. Acta - Mol. Cell Biol. Lipids 1863, 247–265 (2018).

41. Listenberger, L. L. et al. Triglyceride accumulation protects against fatty acid-induced lipotoxicity. Proc. Natl. Acad. Sci. U. S. A. 100, 3077–82 (2003).

42. Zou, Y. et al. A GPX4-dependent cancer cell state underlies the clear-cell morphology and confers sensitivity to ferroptosis. Nat. Commun. 10, 1617 (2019).

43. Yang, W. S. & Stockwell, B. R. Synthetic Lethal Screening Identifies Compounds Activating Iron-Dependent, Nonapoptotic Cell Death in Oncogenic-RAS-Harboring Cancer Cells. Chem. Biol. 15, 234–245 (2008).

44. Wu, J. et al. Intercellular interaction dictates cancer cell ferroptosis via NF2-YAP signalling. Nature 572, 402–406 (2019).

45. Bailey, A. P. et al. Antioxidant Role for Lipid Droplets in a Stem Cell Niche of Drosophila. Cell 163, 340–353 (2015).

46. Ioannou, M. S. et al. Neuron-Astrocyte Metabolic Coupling Protects against Activity-Induced Fatty Acid Toxicity. Cell 177, 1522–1535.e14 (2019).

47. Piskounova, E. et al. Oxidative stress inhibits distant metastasis by human melanoma cells. Nature 527, 186–191 (2015).

48. Stingl, J., Emerman, J. T. & Eaves, C. J. Enzymatic dissociation and culture of normal human mammary tissue to detect progenitor activity. Methods Mol. Biol. 290, 249–63 (2005).

49. Elenbaas, B. et al. Human breast cancer cells generated by oncogenic transformation of primary mammary epithelial cells. Genes Dev. 15, 50–65 (2001).

50. Breunig, C. T. et al. One step generation of customizable gRNA vectors for multiplex CRISPR approaches through string assembly gRNA cloning (STAgR). PLoS One 13, 1–12 (2018).

51. Wang, T., Wei, J. J., Sabatini, D. M. & Lander, E. S. Genetic screens in human cells using the CRISPR-Cas9 system. Science 343, 80–4 (2014).

52. Wiśniewski, J. R., Zougman, A., Nagaraj, N. & Mann, M. Universal sample preparation method for proteome analysis. Nat. Methods 6, 359–362 (2009).

53. Grosche, A. et al. The Proteome of Native Adult Müller Glial Cells From Murine Retina. Mol. Cell. Proteomics 15, 462–480 (2016).

54. van Gorsel, M., Elia, I. & Fendt, S.-M. 13C Tracer Analysis and Metabolomics in 3D Cultured Cancer Cells. Methods Mol. Biol. 1862, 53–66 (2019).

55. Elia, I. et al. Proline metabolism supports metastasis formation and could be inhibited to selectively target metastasizing cancer cells. Nat. Commun. 8, 15267 (2017).

56. Lorendeau, D. et al. Dual loss of succinate dehydrogenase (SDH) and complex I activity is necessary to recapitulate the metabolic phenotype of SDH mutant tumors. Metab. Eng. 43, 187–197 (2017).

57. Fernandez, C. A., Des Rosiers, C., Previs, S. F., David, F. & Brunengraber, H. Correction of 13C Mass Isotopomer Distributions for Natural Stable Isotope Abundance. J. Mass Spectrom. 31, 255–262 (1996).

58. Vriens, K. et al. Evidence for an alternative fatty acid desaturation pathway increasing cancer plasticity. Nature 566, 403–406 (2019).

59. Kharroubi, A. T., Masterson, T. M., Aldaghlas, T. A., Kennedy, K. A. & Kelleher, J. K. Isotopomer spectral analysis of triglyceride fatty acid synthesis in 3T3-L1 cells. Am. J. Physiol. 263, E667–75 (1992).

